# Calorie restriction leads to degradation of mutant uromodulin and ameliorates inflammation and fibrosis in *UMOD*-related kidney disease

**DOI:** 10.64898/2025.12.16.694693

**Authors:** Mariapia Giuditta Cratere, Benedetta Perrone, Barbara Canciani, Céline Schaeffer, Luca Rampoldi

## Abstract

**Background:** Mutations in *UMOD*, encoding uromodulin, lead to Autosomal Dominant Tubulointerstitial Kidney Disease (ADTKD), a genetic cause of kidney failure. *UMOD* mutations have a common gain-of-toxic-function effect, causing mutant uromodulin retention in the endoplasmic reticulum (ER). This leads to ER stress, alteration of protein homeostasis and mitochondrial dynamics, defective autophagy and increased cell death. Calorie restriction (CR) exerts a beneficial role in diseases characterized by accumulation of pathogenic protein and inflammation, by modulating several pathways, including autophagy induction and suppression of inflammation and fibrosis. Given the relevance of these features in ADTKD, we investigated the effect of CR on disease onset and progression.

**Methods:** Transgenic mice expressing C147W uromodulin (Tg*^Umod^*^C147W^) were subjected to a moderate (30%) CR regimen for 15 or 24 weeks, starting at different stages of disease progression.

**Results:** CR restored autophagy, as shown by decreased P62 punctae and quenched mTOR activation specifically in mutant uromodulin expressing cells, it recovered expression of key ER-phagy receptor genes, with a concomitant, striking reduction of mutant uromodulin ER retention. In pre-symptomatic Tg*^Umod^*^C147W^ mice, CR alleviates epithelial cell stress. This, likely along with a direct anti-inflammatory effect of CR, prevents inflammation and progressive decline of kidney function. At this early disease stage, CR ameliorates the already established kidney damage and reduces fibrosis, suggesting reversal of ADTKD phenotype. CR was also effective in significantly delaying disease progression in Tg*^Umod^*^C147W^ mice with advanced disease and already compromised kidney function.

**Conclusion:** Our findings establish the proof-of-principle that counteracting the primary effect of *UMOD* mutations by stimulating autophagy and quenching inflammation prevents disease onset and progression. This study uncovers the potential of CR as a valuable therapeutic option in the context of ADTKD-*UMOD*, and possibly other proteinopathies.

**Key points:**

- Calorie restriction (CR) stimulates autophagy to degrade mutant uromodulin leading to amelioration of cell stress and tubular damage.
- At early disease stage CR reverts ADTKD-*UMOD* phenotype, by preventing inflammation and progressive decline of kidney function and rescuing fibrosis.
- At advanced disease stage, CR significantly delays disease progression and worsening of kidney function.

## Introduction

*UMOD*-related Autosomal Dominant Tubulointerstitial Kidney Disease (ADTKD-*UMOD*; MIM #16200) is a Mendelian disorder characterized by interstitial fibrosis with diffuse tubulointerstitial damage, urine concentration defect, hyperuricemia and gout. It causes progressive loss of kidney function leading to kidney failure, predominantly between the third and sixth decade of life.^1,2^ ADTKD-*UMOD* is caused by mutations in the *UMOD* gene^3^ and represents the third most frequent kidney monogenic disorder^4^ accounting for 2% of all kidney failure cases.^5^

*UMOD* encodes uromodulin^6,7^, the major urinary protein that is specifically produced in the kidney by tubular epithelial cells (TECs) of the thick ascending limb of Henle’s loop (TAL) and of the early distal convoluted tubules.^8^ *UMOD* mutations are predominantly missense^9,10^ and exert a shared cellular effect. They all affect protein folding and trafficking, causing misfolded uromodulin to accumulate in the endoplasmic reticulum (ER) of TECs, with subsequent induction of ER stress and the unfolded protein response. These features are observed in patient samples^11–14^ and recapitulated in several *in vitro*^15–18^ and *in vivo* models^11,14,19–22^, ascribing ADTKD-*UMOD* as an ER storage disease^23^. The different extent of ER retention induced by *UMOD* mutations associates with disease severity in patients, and it was proposed as a predictor of age at kidney failure.^9^ Besides ER retention, gene dosage and propensity of mutant uromodulin to aggregate has been recently demonstrated to play a key role in disease pathogenesis and it is directly correlated with disease severity in patients and corresponding mouse models.^14^ Secondary phenotypes of the disease, likely downstream of ER stress, are altered mitochondria homeostasis and dynamics^21,22^, defective protein homeostasis, increased apoptosis ^19^ and impaired autophagy in TAL cells.^14,19,22^ Inflammation is a key disease feature and an early event of the pathogenetic cascade, preceding fibrosis and occurring concomitantly with TAL cell stress and tubular damage.^24^

The mechanisms linking mutant uromodulin ER retention to kidney damage or ways to counteract the effects of uromodulin mutations are only partially clarified. In fact, an effective, specific therapeutic strategy is presently an unmet clinical need.

Given the gain-of-(toxic)-function effect of *UMOD* mutations, removal of intracellularly retained mutant protein could be a valuable option. This is supported by recent evidence showing that reducing mutant uromodulin intracellular accumulation, by promoting its trafficking and secretion, led to an improvement of kidney damage in an *in vivo* model of the disease.^25^ Similarly, increasing autophagy might be advantageous, as suggested by recent data showing that stimulation of autophagy leads to degradation of mutant uromodulin aggregates in cell models^14^ and that overexpression of the mesencephalic astrocyte derived neurotrophic factor (MANF), a proposed autophagy inducer, ameliorates kidney phenotype in an ADTKD-*UMOD* mouse model.^22^

Calorie restriction (CR), consisting in reducing daily calorie intake by 20 to 50% without causing malnutrition^26^ was reported to be beneficial in different disease settings characterized by accumulation of pathogenic proteins and inflammation.^27,28^ This protective role of CR is mediated by its action on several molecular mechanisms, including improvement of mitochondrial and autophagy dysfunction and suppression of inflammation and oxidative stress.^29^

Considering that all these are key features of the disease molecular pathogenesis^14,19,21,22,24^, here we investigated the therapeutic potential of CR in ADTKD-*UMOD.* To this end, transgenic mice expressing C147W mutant uromodulin and recapitulating the human disease^11,24^ were subjected to a moderate 30% CR regimen^30,31^ for 15 or 24 weeks starting at different stages of disease progression. At early stage of the disease CR restored autophagy in uromodulin expressing cells and induced degradation of mutant uromodulin, leading to a striking reduction of ER retained protein. Moreover, CR attenuated TECs stress and reduced inflammation and subsequent fibrosis, eventually preventing decline of kidney function and reverting the already established kidney damage. Importantly, CR was effective in counteracting disease progression also when started in mice with advanced disease and compromised kidney function.

Our results point at CR, or possibly other autophagy-inducing interventions, as a potential and effective therapeutic option in the context of ADTKD-*UMOD* and establish the proof of principle that counteracting the primary effect of *UMOD* mutations can prevent disease onset and progression.

## Methods

### Animal models

All animals used in this study were housed under specific pathogen-free conditions with controlled humidity and temperature on a 12-h dark/light cycle and *ad libitum* (AL) access to tap water and standard chow (Diet VRF1 (P), SDS Diet, Augy, FR), except during periods of CR experiments (see below). All procedures were reviewed and approved by the Institutional Animal Care and Use Committee (IACUC) of the San Raffaele Institute and were performed in accordance with the institutional guidelines (protocol #1368, #1526). Transgenic mouse lines expressing mutant (Tg*^Umod^*^C147W^) or wild-type (Tg*^Umod^*^wt^) uromodulin (i.e., transgenic mice for wild-type uromodulin, serving as an expression matched control), both on the FVB/NJ background, were previously generated and characterized.^11,24^ Sequences of primer sets used for genotyping are listed in **Supplemental Table 1**.

### CR protocols

To test the effect of 30% CR on disease at early stages, a 15-week-long CR experiment was conducted on Tg*^Umod^*^C147W^ mice starting at 8 weeks of age. To assess the effect of 30% CR at advanced stage of disease 24-week-old Tg*^Umod^*^C147W^ mice were treated for 24 weeks. For each CR protocol experimental animals were randomly divided into AL or CR groups at the designated timepoint and individually housed to measure the exact food intake. Food consumption of the animals in the AL groups was determined by weighting leftover chow daily. Each day animals in the CR groups were allotted 70% of the food consumed by the mice in the AL regimen, as previously reported.^32–34^ Body weight of animals was assessed weekly. For all experiments, age-matched male and female mice were used. The number of mice for each experiment is reported in figure legends. Data obtained shown in graphs as squares for male mice and as circles for female mice. All data for each experimental group are shown including both sexes, except for 15-week-old CR protocol, for which data for male mice are reported in **Supplemental Figure 1**.

### Statistical analysis

Data are reported as means ± standard deviation (s.d.) or standard error of mean (s.e.m.). Data statistical analysis was performed with Prism V10.4.2 software (GraphPad Inc.). Unpaired Student’s t-test was performed when comparing mice of the same genotype subjected to different diets. Comparison between more than two experimental groups was performed by using one or two-way ANOVA followed by Dunnett’s, Šídák’s or Tukey’s correction as indicated in figure legends. A *P*<0.05 was considered statistically significant. Multivariate data analysis was performed using unsupervised Principal Component Analysis (PCA), to integrate all parameters tested to assess the effect of CR at early and late disease stage. Since such parameters were measured according to different scales, data were normalized by auto-scaling transformation to ensure comparability across variables. Experimental groups were visualized with 95% confidence interval ellipses that were automatically generated by the software to indicate group clustering. To evaluate the statistical significance of group separation, Permutational Multivariate Analysis of Variance (PERMANOVA) was performed. This method considers: 1) the F-value, which compares intra- and inter-group variance; 2) the R-squared value, representing the proportion of variation in the distance matrix explained by the grouping factor (0, no variation explained; 1, all the variation explained), and 3) the *P*-value based on 999 permutations. PCA, heatmaps of disease parameters and K-means clustering analyses were generated by using MetaboAnalyst 6.0. online package (www.metaboanalyst.ca/home.xhtml).

### Additional methods

Detailed methods are provided in **Supplemental Methods** and **Supplemental Tables 1**, **10** and **11**.

## Results

### CR reduces mutant uromodulin ER retention and aggregation

To test the hypothesis that reducing calorie intake could lead to reduction of mutant uromodulin intracellular load, we performed a 15-week-long, 30% CR experiment on Tg*^Umod^*^C147W^ and Tg*^Umod^*^wt^ mice^11^. The initial time point (8 weeks) corresponds to an early phase of disease, with signs of kidney inflammation and fibrosis but preserved kidney function.^24^ At the final time point (23 weeks), Tg*^Umod^*^C147W^ mice show the full-blown disease, with declined kidney function^11^ (**Figure 1A, Supplemental Figure 1A**). Body weight of mice under CR decreased (<20% of the initial body weight) during in the first 3 weeks of diet and it then stabilized until the end of treatment (**Figure 1B, C** and **Supplemental Figure 1B, C**). As compared to Tg*^Umod^*^wt^ mice, AL Tg*^Umod^*^C147W^ mice showed uromodulin accumulation within the kidney, that is massively decreased upon CR (**Figure 1D** and **Supplemental Figure 1D**). Such reduction is mostly contributed by a significant decrease of uromodulin ER precursor (90kDa band, corresponding to immature uromodulin carrying ER-type glycans)^11^, reflecting reduced ER retention of mutant protein after CR (**Figure 1D** and **Supplemental Figure 1D**). This is confirmed by immunofluorescence staining, specifically analysing the distribution of transgenic uromodulin and showing decreased colocalization of mutant protein with the ER chaperone calreticulin (CLR) (**Figure 1E** and **Supplemental Figure 1E**). Such effect is associated with a strong decrease of mutant uromodulin aggregates, as shown by decreased level of high molecular weight forms of uromodulin on Western blot in non-reducing conditions^14^ (**Figure 1F** and **Supplemental Figure 1F**). As previously reported and consistent with impaired protein maturation, AL Tg*^Umod^*^C147W^ mice showed striking reduction of urinary uromodulin levels. After CR, reduction of uromodulin levels in the kidney was not accompanied by increased urinary excretion (**Figure 1G** and **Supplemental Figure 1G**). The effect of the CR protocol is not due to changes of gene expression, as shown by comparable transcript levels of total and transgenic *Umod* in all experimental groups (**Figure 1H** and **Supplemental Figure 1H**). Altogether these results demonstrate that CR is effective in reducing mutant uromodulin ER retention and aggregation, without altering its expression, trafficking or urinary secretion.

**Figure 1.**
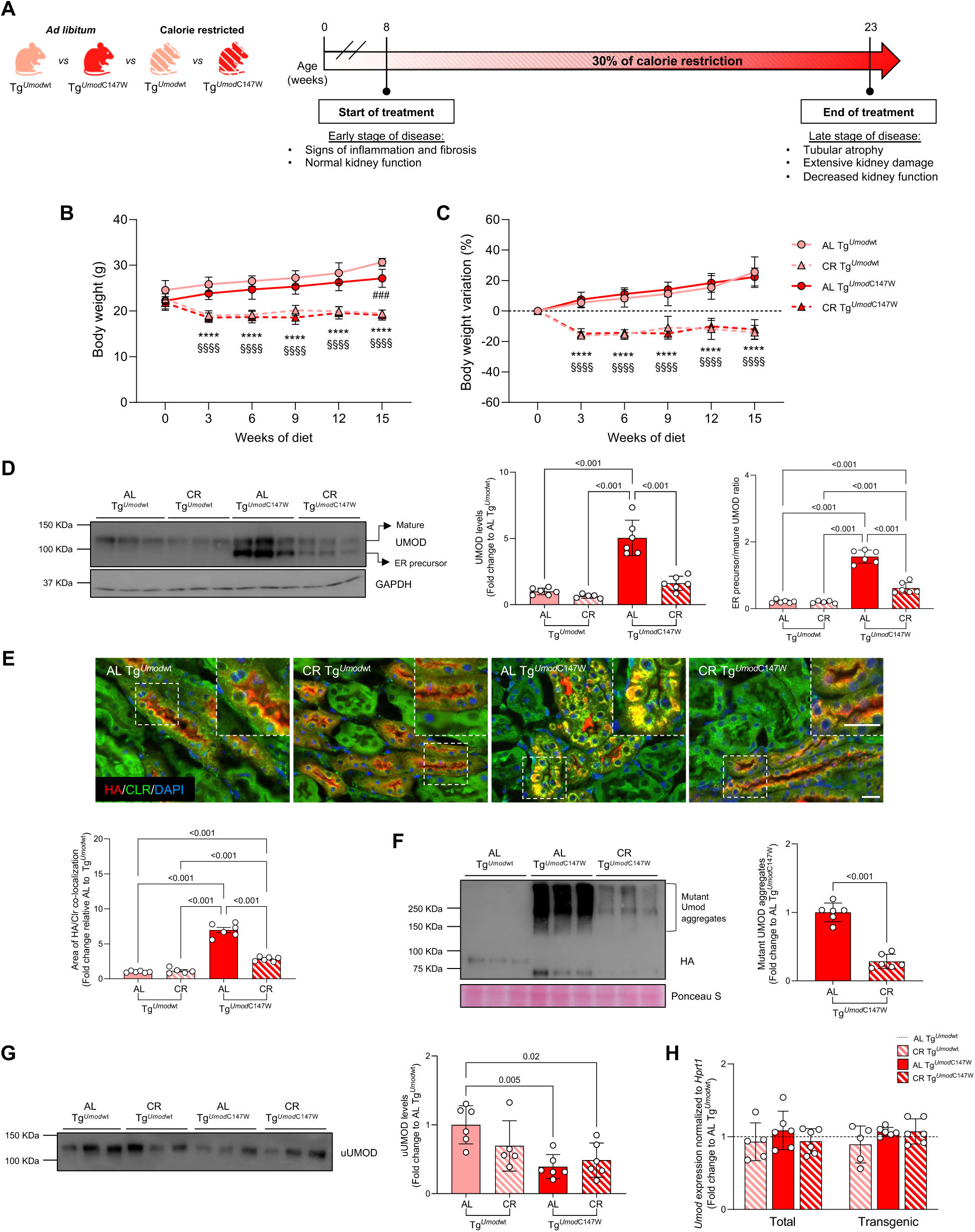
CR reduces mutant uromodulin ER retention and aggregation. **A**) Graphical representation of the CR protocol. **B, C**) Body weight (**B**) and percentage of weight loss compared to the initial timepoint (**C**) of mice during the diet. ^###^*P*<0.001 AL Tg*^Umod^*^wt^ vs AL _Tg_*_Umod_*_C147W; \*\*\*\**P*<0.0001 AL Tg_*_Umod_*_wt vs CR Tg_*_Umod_*_wt; §§§§*P*<0.0001 AL Tg_*_Umod_*_C147W vs CR_ Tg*^Umod^*^C147W^. **D**) Western blot analysis of UMOD in kidney lysates. The ER precursor (immature, 90 kDa) and mature form of UMOD (120 kDa) are indicated. GAPDH is reported as a normalizer. Graphs represent quantification of total UMOD kidney levels and the ratio of ER precursor to mature UMOD isoforms. **E**) Immunofluorescence analysis of transgenic UMOD (HA, in red) and the ER marker calreticulin (CLR, in green). Nuclei are counterstained with DAPI (blue). The magnified field is indicated by a dashed square on each picture. Scale bar 20 μm. Histograms represent the quantification of the area of transgenic UMOD-CLR co-localization. **F**) Western blot in non-reducing conditions of transgenic UMOD (anti-HA). High molecular-weight (HMW) signal corresponds to mutant UMOD aggregates. Ponceau S staining is shown as a loading control. Histograms represent the quantification of UMOD aggregates in kidneys from AL or CR Tg*^Umod^*^C147W^ mice. **G**) Western blot and quantification of urinary UMOD (uUMOD) levels. Urine samples were normalized over total volume of 16-hour urine collection in metabolic cage. **H**) Total and transgenic *Umod* transcript levels measured by real-time RT-qPCR in kidneys from AL and CR Tg*^Umod^*^wt^ and Tg*^Umod^*^C147W^ mice. n=5-6 mice/group (all females). Data are reported as fold change relative to AL Tg*^Umod^*^wt^ (in **D, E, G** and **H**) or AL Tg*^Umod^*^C147W^ (in **F**). Bars represent mean ± s.d. (in **B, C, D, F, G** and **H**) or mean ± s.e.m. (in **E**). Group comparisons were performed by one-way ANOVA followed by followed by Šidák’s (in **B** and **C**) or Tukey’s (in **D, E** and **G**) corrections or unpaired t-test (in **F**).

### CR ameliorates ADTKD-*UMOD* kidney phenotype

We then analysed inflammation and fibrosis, two important ADTKD features. All histological parameters of kidney damage (tubular dilation, inflammatory cell infiltration and interstitial fibrosis) were increased in AL Tg*^Umod^*^C147W^ mice, as expected, and significantly reduced to levels comparable to Tg*^Umod^*^wt^ micev upon CR (**Figure 2A-D** and **Supplemental Figure 2A, B**). A similar trend was observed for the renal expression of inflammation and fibrosis related genes. They were all upregulated in AL Tg*^Umod^*^C147W^ mice and remarkably rescued by CR (**Figure 2E, Supplemental Table 2** and **Supplemental Figure 2C**). CR reduced the expression of some genes related to inflammation (i.e., *Ptprc*, *Ccl5*) also in kidneys of Tg*^Umod^*^wt^ mice relative to their AL counterpart, substantiating the already described impact of CR on inflammatory cell metabolism.^35,36^

**Figure 2.**
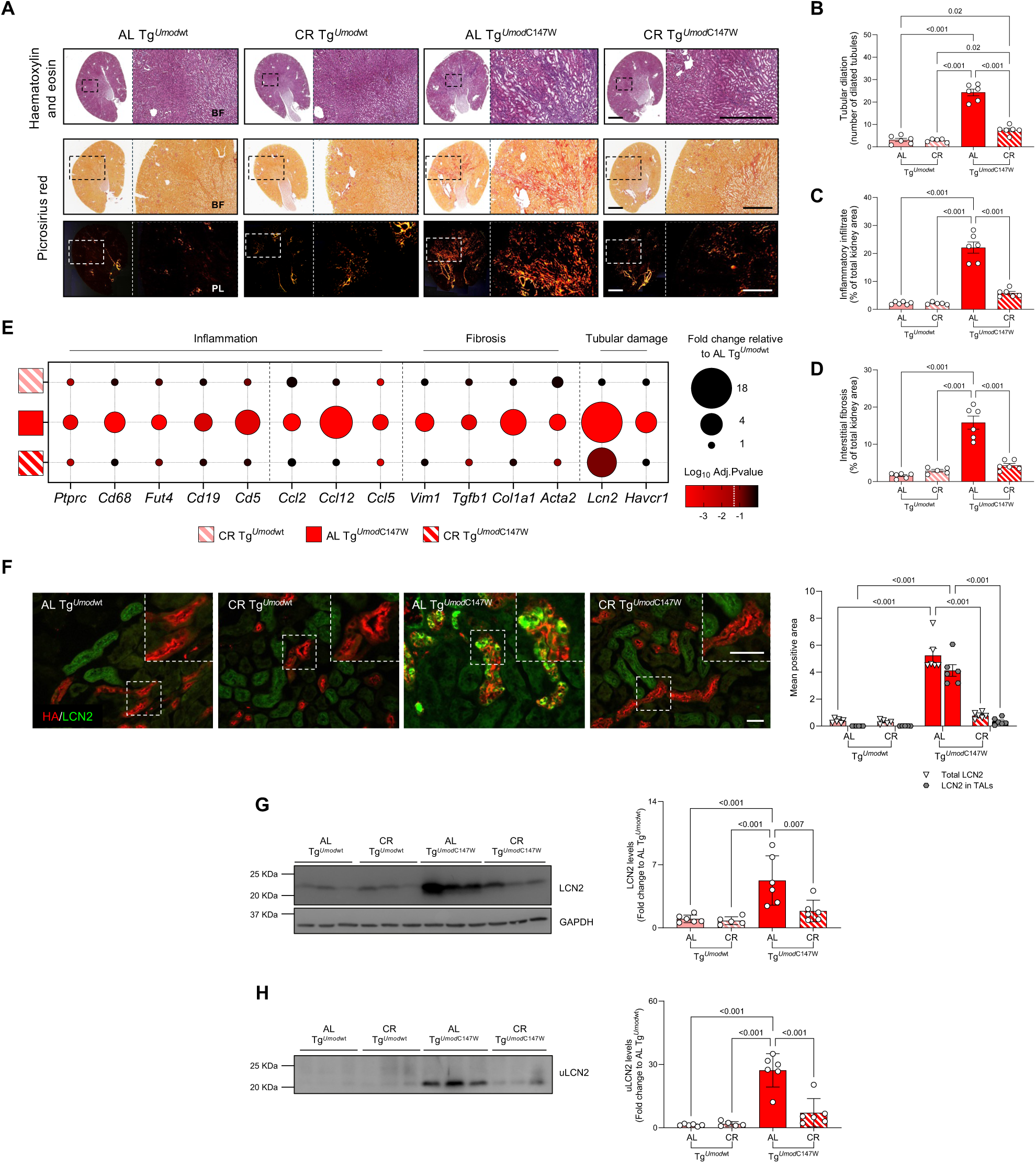
CR ameliorates kidney inflammation, fibrosis and tubular damage. **A**) Representative histology images of kidney sections stained with haematoxylin and eosin or picrosirius red and viewed in bright field (BF) or under polarized light (PL). Scale bars 2 mm and 600 μm. **B, C, D**) Graphs represent quantification of tubular dilation (**B**) and areas of inflammatory cell infiltration (**C**) and interstitial fibrosis (**D**), expressed as percentage of total kidney area. **E**) Categorical bubble plot showing the kidney expression levels of genes encoding for inflammatory cell markers (*Ptprc*, leucocytes; *Cd68*, macrophages, *Fut4*, neutrophils; *Cd19*, B-cells; *Cd5*, T-cells), chemokines (*Ccl2*, *Ccl12*, *Ccl5*), fibrosis markers (*Vim1*, *Tgfb1*, *Col1a1*, *Acta2*) and markers of tubular damage (*Lcn2* and *Havcr1*). Transcript levels were analysed by real-time RT-qPCR. Bubble size is proportional to fold change relative to AL Tg*^Umod^*^wt^ mice. Bubble colour corresponds to Log_10_ of the adjusted *P* value relative to AL Tg*^Umod^*^wt^ (one-way ANOVA followed by Dunnett’s multiple comparisons post-hoc test). A Log_10_ adjusted *P* value ≤ −1.3 (white dashed line) was considered statistically significant. Data used to generate the categorical bubble plot are reported in *Supplemental Table 2*. **F**) Immunofluorescence staining for transgenic UMOD (HA, in red) and LCN2 (in green) in kidney sections from AL or CR Tg*^Umod^*^wt^ and Tg*^Umod^*^C147W^ mice. The magnified field is indicated by a dashed square on each picture. Scale bar 40 μm. Histograms represent quantification of the LCN2 positive signal in total kidney area or in TAL tubules (HA^+^). **G, H**) Western blot analysis and quantification of LCN2 levels in kidney (**G**) and urine (uLCN2) (**H**). For kidney lysates, GAPDH was used as a loading control. Urine samples were normalized over total urine volume collected in 16 hours. n=5-6 mice/group (all females). Bars represent mean ± s.e.m. (in **B, C, D** and **F**) or ± s.d. (in **G** and **H**). In **G** and **H**, data are reported as fold change relative to AL Tg*^Umod^*^wt^. Group comparisons were performed by one-way (in **B, C, D, G** and **H**) or two-way ANOVA (in **F**) followed by Tukey’s correction.

Tubular damage is an early event occurring in mutant uromodulin expressing mice. Lipocalin-2 (Lcn2), a protein expressed in injured TECs^37^ is already induced in kidneys of mice at 1 week of age. Its expression remains high in adult mice, and it represents one of the most up-regulated genes identified by kidney transcriptional profiling.^14,24^ As the disease progresses, we also reported induction of *Havcr1* (encoding kidney injury molecule-1, Kim1)^38^ starting at 4 weeks of age.^24^ Upon CR, *Lcn2* and *Havcr1* expression levels were significantly reduced in kidneys of Tg*^Umod^*^C147W^ mice compared with their AL counterpart (**Figure 2E**). Immunofluorescence analysis revealed that LCN2 signal is specifically induced in mutant uromodulin expressing cells, and it is dramatically reduced, along with mutant protein aggregates, in CR Tg*^Umod^*^C147W^ mice (**Figure 2F**). This is reflected by significant decrease of LCN2 protein levels in kidneys (**Figure 2G**) and urines (**Figure 2H**), implying that reducing mutant uromodulin ER retention and aggregation can rescue TAL cell damage in Tg*^Umod^*^C147W^ mice.

These results demonstrate that CR effectively ameliorates ADTKD-*UMOD* phenotype, by reducing inflammation and fibrosis and rescuing tubular damage.

### CR restores autophagy in mutant uromodulin expressing cells

Since reduction of mutant uromodulin accumulation in kidney lysates of CR mice is not reflected by reduced gene expression nor by increased secretion, we hypothesized that this could be explained by protein degradation. To gain mechanistic insight, we evaluated markers of autophagy. Induction of this pathway was recently demonstrated to play a role in degrading mutant uromodulin aggregates in cell models^14^ and to have a beneficial effect on disease phenotype *in vivo*.^22^ Moreover, CR was reported to stimulate autophagy by suppressing the activity of the mammalian target of rapamycin (mTOR) signalling pathway, regulating cell catabolism.^39^

We tested the effect of CR on the expression of P62, an autophagy receptor that is itself degraded along with the substrate, thus representing a readout of autophagic flux.^40^ In kidneys of AL Tg*^Umod^*^C147W^ mice we observed increased P62 protein levels, and specific accumulation of P62-positive punctae in cells expressing mutant uromodulin (**Figure 3A, B**). This was associated with comparable levels of transcript (*Sqstm1*) in kidneys of mutant mice compared with Tg*^Umod^*^wt^ mice (**Figure 3C**). Such effect can hence be considered evidence of reduced/impaired autophagic flux^41^ specifically in mutant uromodulin expressing cells, consistent with results in other ADTKD-*UMOD* models.^14,19,22^ Noteworthy, accumulation of P62 punctae as well as its kidney levels were strongly reduced in TALs of CR Tg*^Umod^*^C147W^ mice (**Figure 3A, B**), despite stable transcript levels (**Figure 3C**), suggesting CR-mediated rescue of autophagic flux. This was also demonstrated by kidney levels of LC3-II/LC3-I ratio, that are increased in AL Tg*^Umod^*^C147W^ mice and reduced to levels comparable to Tg*^Umod^*^wt^ mice upon CR (**Figure 3D**).

**Figure 3.**
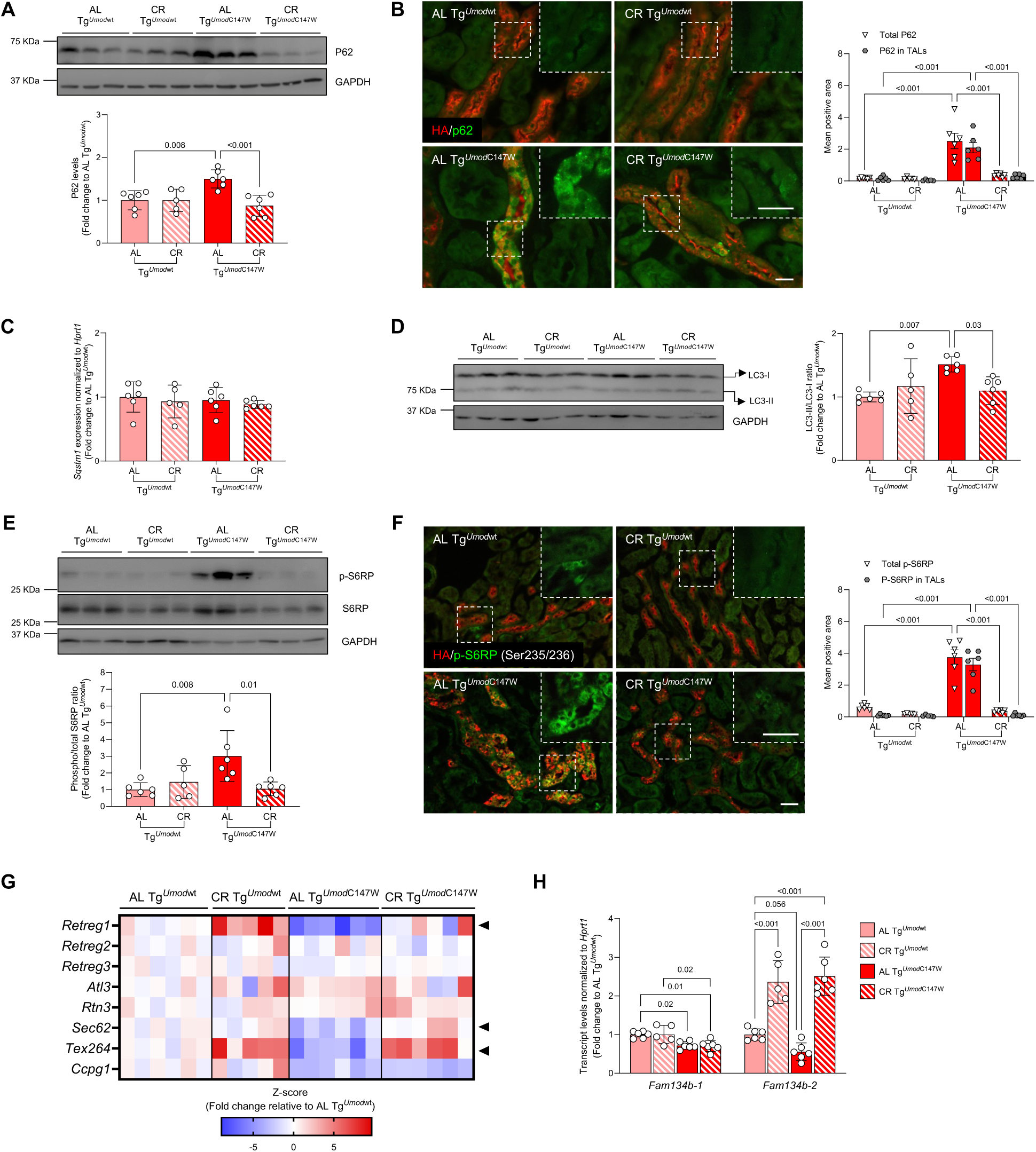
CR restores autophagy in mutant uromodulin expressing cells. **A**) Western blot analysis and quantification of P62 kidney levels. GAPDH is reported as a loading control. **B**) Immunofluorescence staining for P62 (in green) and transgenic UMOD (HA, in red) in kidney sections. The magnified field is indicated by a dashed square on each picture. Scale bar 20 μm. Histograms represent quantification of the P62 positive signal in total kidney area or in TAL tubules (HA^+^). **C**) Analysis of *Sqstm1* (encoding P62) kidney expression levels. **D**) Western blot analysis of LC3 (autophagosome marker) and quantification of LC3-II/LC3-I (active/inactive) ratio. **E**) Western blot analysis of phosphorylated (p-S6RP; Ser235/236) and total S6 ribosomal protein (S6RP; downstream target of mTOR activity) in total kidney lysates. The graph shows the phosphorylated/total S6RP ratio. GAPDH is reported as a loading control. **F**) Immunofluorescence staining for transgenic UMOD (HA, in red) and p-S6RP (in green) in kidney sections. The magnified field is indicated by a dashed square on each picture. Scale bar 20 μm. Histograms represent quantification of the p-S6RP positive signal in total kidney or in TAL tubules (HA^+^). **G**) Heatmap showing the kidney expression levels of ER-phagy receptor genes *Retreg1* (encoding Fam134b), *Retreg2* (Fam134a), *Retreg3* (Fam134c), *Atl3*, *Rtn3*, *Sec62*, *Tex264*, *Ccpg1* in AL or CR Tg*^Umod^*^wt^ and Tg*^Umod^*^C147W^ mice. Transcript levels were analyzed by real-time RT-qPCR. For each gene, expression levels are reported as z-score of the fold change relative to AL Tg*^Umod^*^wt^. Arrow heads indicate genes that are significantly downregulated in kidneys of AL Tg*^Umod^*^C147W^ mice compared to AL Tg*^Umod^*^wt^ mice and whose expression level is restored upon CR. For all the analyzed genes, the fold change relative to AL Tg*^Umod^*^wt^ mice and statistical analysis is reported in *Supplemental Table 3*. **H**) Transcript levels of *Fam134b-1* (full-length) and *Fam134b-2* (N-terminal truncated, starvation-induced *Fam134b* isoform) in kidneys of AL or CR Tg*^Umod^*^wt^ and Tg*^Umod^*^C147W^ mice. n=5-6 mice/group (all females). Bars represent mean ± s.d. (in **A, C, D, E** and **H**) or ± s.e.m. (in **B** and **F**). In **A, C, D, E** and **H** data are reported as fold change relative to AL Tg*^Umod^*^wt^. Group comparisons were performed by one-way (in **A, D, E** and **H**) or two-way (in **B** and **F**) ANOVA followed by Tukey’s correction.

To assess the potential involvement of mTOR signalling pathway in CR-induced autophagy, we measured phosphorylation of its downstream target, ribosomal protein SR (S6RP). Compared to Tg*^Umod^*^wt^ mice, AL Tg*^Umod^*^C147W^ mice show increased kidney levels of phospho-S6RP that are almost exclusively observed in mutant uromodulin expressing cells. Notably, the levels of S6RP were essentially normalised by CR (**Figure 3E, F**).

Given the reduction of ER-retained mutant uromodulin in TAL cells upon CR, we investigated ER-phagy, a form of autophagy mediated by selective receptors, that specifically degrades portions of the ER.^42^ We tested the kidney expression levels of genes encoding for some of the ER-phagy receptors so far reported in mammals.^43^ Among these receptors *Retreg1*, *Sec62* and *Tex264* appeared to be significantly decreased in kidneys of mutant mice in basal conditions and their expression was restored to Tg*^Umod^*^wt^ levels upon CR (**Figure 3G** and **Supplemental Table 3**). Since an N-terminal truncated isoform of FAM134B (FAM134B-2) encoded by *Retreg1* has been demonstrated to be transcriptionally enhanced by starvation regulating starvation-induced ER-phagy^44–46^, we specifically analysed the expression of the different FAM134B isoforms. We demonstrate that *Fam134b-1* (encoding the full-length protein) and *Fam134b-2* are down-regulated in kidneys of AL Tg*^Umod^*^C147W^ mice and only *Fam134b-2* is up-regulated in kidneys of CR Tg*^Umod^*^C147W^ mice (**Figure 3H**), suggesting a possible role of CR in recovering ER-phagy impairment in kidneys of Tg*^Umod^*^C147W^ mice.

These data strongly suggest that, by inhibiting mTOR overactivation in kidneys of mice expressing mutant uromodulin, CR rescues autophagy (and likely ER-phagy) in TAL cells, thus allowing degradation of ER-retained mutant uromodulin.

### CR reverts ADTKD phenotype at early disease stage

Next, we assessed the impact of CR on disease progression by comparing 23-week-old AL and CR Tg*^Umod^*^C147W^ mice with 8-week-old AL Tg*^Umod^*^C147W^ mice, corresponding to the age of animals at the beginning of treatment (**Figure 4A**). To evaluate kidney function, we performed plasma and urine biochemistry at the beginning and at the end of dietary intervention. At 23 weeks of age, AL Tg*^Umod^*^C147W^ mice developed significant polyuria and signs of kidney failure, as demonstrated by progressive rise of blood urea nitrogen (BUN) and diuresis levels (**Figure 4B, C** and **Supplemental Table 4, 5**). Both these parameters remained stable and within the normal range in CR Tg*^Umod^*^C147W^ mice. Importantly, variation in these parameters is not an effect of CR *per se*, since they show no significant change in CR Tg*^Umod^*^wt^ mice compared to their AL counterpart (**Supplemental Table 4, 5**). These results demonstrate that CR prevents decline of kidney function in Tg*^Umod^*^C147W^ mice. ER accumulation of mutant uromodulin is already evident in TALs of 8-week-old Tg*^Umod^*^C147W^ mice (**Figure 4D-F**), and the extent of ER retention significantly increases in 23-week-old AL Tg*^Umod^*^C147W^ mice, indicating disease progression. In contrast, in kidneys of 23-week-old CR Tg*^Umod^*^C147W^ mice, mutant uromodulin aggregates appeared reduced to levels even lower than those measured at 8 weeks of age (**Figure 4D-F**). Such reduction is not due to *Umod* expression, which is in fact increased with age in kidneys of Tg*^Umod^*^C147W^ mice, irrespective of their dietary regimen (**Figure 4G**). These data demonstrate that CR not only prevents mutant uromodulin accumulation but is also effective in promoting clearance of existing protein aggregates.

**Figure 4.**
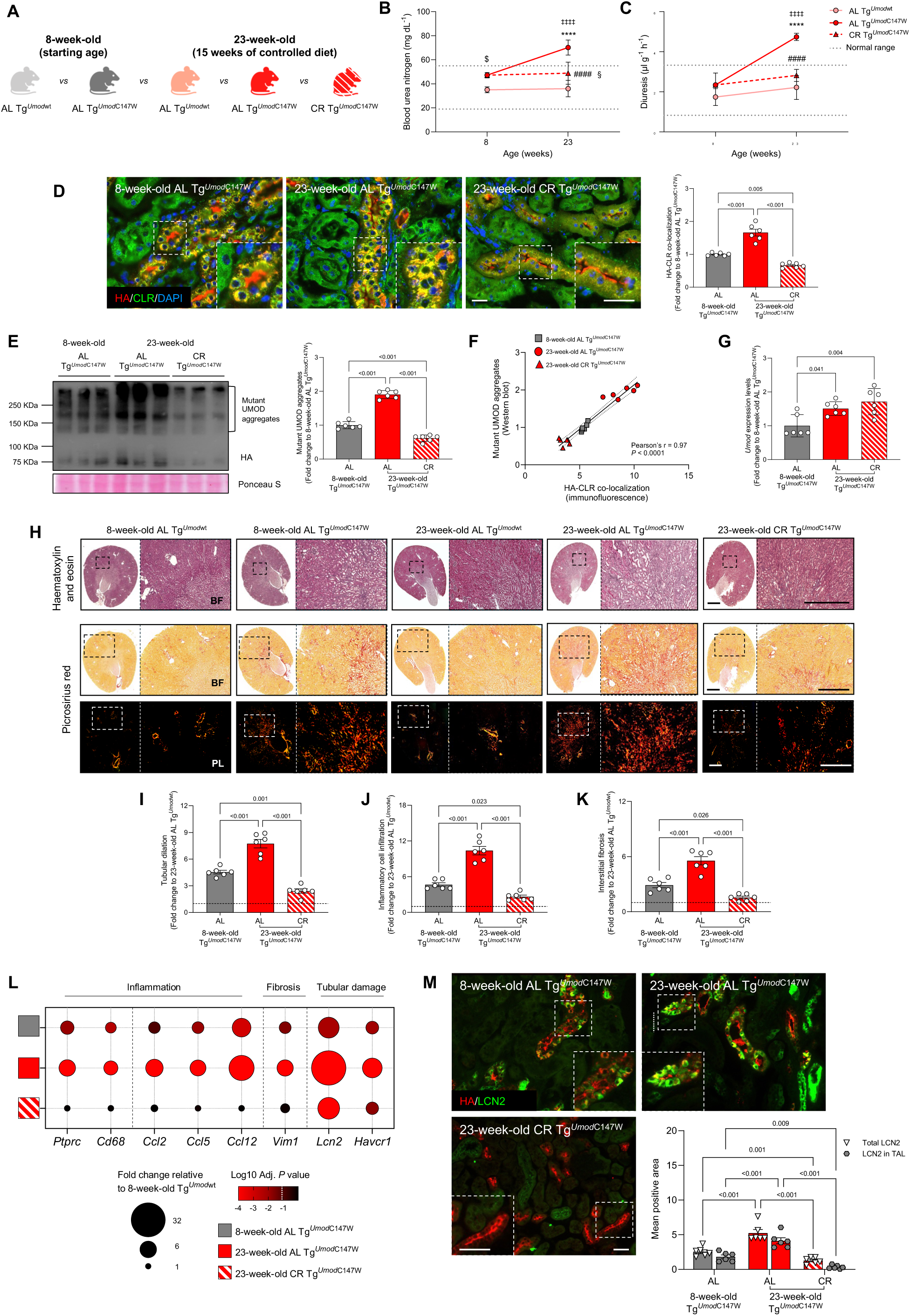
CR prevents ADTKD progression and reverts disease phenotype at early disease stage. **A**) To assess the potential of CR on ADTKD-*UMOD* phenotype reversal, 23-week-old AL Tg*^Umod^*^wt^ mice, AL or CR Tg*^Umod^*^C147W^ mice were compared with 8-week-old AL Tg*^Umod^*^wt^ and Tg*^Umod^*^C147W^ mice, corresponding to the age of mice at the beginning of the dietary intervention. **B, C**) BUN (**B**) and diuresis (**C**) levels of mice at 8 weeks and 23 weeks of age. Grey dashed lines represent normal range values. ^$^*P*<0.05 8-week-old AL Tg*^Umod^*^wt^ vs 8-week-old AL Tg*^Umod^*^C147W^; \*\*\*\**P*<0.0001 23-week-old AL Tg*^Umod^*^wt^ vs 23-week-old AL Tg*^Umod^*^C147W^; ^####^*P*<0.0001 23-week-old AL Tg*^Umod^*^C147W^ vs 23-week-old CR Tg*^Umod^*^C147W^; ^§^*P*<0.05 23-week-old AL Tg*^Umod^*^wt^ vs 23-week-old CR Tg*^Umod^*^C147W^; ^‡‡‡‡^*P*<0.0001 8-week-old AL Tg*^Umod^*^C147W^ vs 23-week-old AL Tg*^Umod^*^C147W^. **D**) Immunofluorescence staining for transgenic UMOD (HA, in red) and CLR (in green). Nuclei are counterstained with DAPI (in blue). Scale bar 40 μm. Histograms represent quantification of transgenic UMOD-CLR co-localization area. **E**) Western blot in non-reducing conditions to detect transgenic UMOD (HA) in kidney lysates. Ponceau S staining is reported as a loading control. Graph represents quantification of HMW mutant UMOD aggregates. **F**) Linear regression illustrating the correlation between mutant UMOD aggregates measured by Western blot and HA-CLR signal co-localization measured by immunofluorescence. The dotted lines show the 95% confidence intervals. Dots represent individual animals. **G**) *Umod* transcript levels measured by real-time RT-qPCR. **H**) Representative histology images of kidney sections stained with haematoxylin and eosin or picrosirius red and viewed in bright field (BF, upper panels) or under polarized light (PL, bottom panels). Scale bars 2 mm and 600 μm. **I, J, K**) Quantification of tubular dilation (**I**), inflammatory cell infiltrate (**J**) and interstitial fibrosis (**K**). **L**) Categorical bubble plot showing the kidney expression levels of inflammation (*Ptprc*, *Cd68*, *Ccl2*, *Ccl5*, *Ccl12*), fibrosis (*Vim1*) and tubular damage (*Lcn2* and *Havcr1*) related genes. Transcript levels were analysed by real-time RT-qPCR. Bubble size is proportional to fold change relative to 8-week-old AL Tg*^Umod^*^wt^. Bubble colour corresponds to Log_10_ of the adjusted P value relative to 8-week-old AL Tg*^Umod^*^wt^ mice (one-way ANOVA followed by Dunnett’s multiple comparisons post-hoc test). A Log_10_ adjusted *P* value ≤ −1.3 (white dashed line) was considered statistically significant. Data used to generate the categorical bubble plot are reported in *Supplemental Table 6*. **M**) Representative immunofluorescence staining for transgenic UMOD (HA, in red) and LCN2 (in green) in kidney sections. The magnified field is indicated by a dashed square on each picture. Scale bar 40 μm. Histograms represent quantification of the LCN2 positive signal in total kidney area or in TAL tubules (HA^+^). Bars represent mean ± s.d. (in **B, C, E, G**) or ± s.e.m. (in **D, I, J, K, M**). Data are reported as fold change relative to 8-week-old AL Tg*^Umod^*^C147W^ (in **D, E, G**) or to 8-week-old AL Tg*^Umod^*^wt^ (dashed line in **I, J, K**). Group comparisons were performed by one-way ANOVA followed by Šidák’s (in **B** and **C**) or Tukey’s (in **D, E, G, I, J** and **K**) correction, or two-way ANOVA followed by Tukey’s correction (in **M**).

In Tg*^Umod^*^C147W^ mice disease progression is also evidenced by progressive increase of tubular dilation, inflammation and interstitial fibrosis (**Figure 4H-L**, **Supplemental Figure 3A, B** and **Supplemental Table 6**) and in the expression of markers of tubular damage, such as *Lcn2* (**Figure 4L, M** and **Supplemental Figure 3C**) and *Kim1* (**Figure 4L**). Notably, upon CR, tubular dilation and levels of macrophage infiltration and extracellular matrix deposition in kidneys of 23-weeks-old Tg*^Umod^*^C147W^ mice were comparable to what measured in wild-type animals and appeared significantly decreased compared to 8-week-old AL Tg*^Umod^*^C147W^ mice (**Figure 4H-L**, **Supplemental Figure 3A, B** and **Supplemental Table 6**). Moreover, CR strongly attenuates tubular damage in kidneys of mutant mice, as demonstrated by significant reduction of LCN2 in TAL tubules of 23-week-old CR Tg*^Umod^*^C147W^ mice, compared to their genotype-matched counterpart at 8 weeks of age (**Figure 4M** and **Supplemental Figure 3C**).

These data collectively demonstrate that, when started at early stage of disease, CR prevents tubular damage and disease progression and rescues the already present kidney damage in Tg*^Umod^*^C147W^ mice.

### CR blocks disease progression in mice with advanced ADTKD-*UMOD*

To substantiate the translational relevance of CR, we then investigated its effectiveness at an advanced disease stage. Tg*^Umod^*^C147W^ mice were fed a similar CR regimen starting at 24 weeks of age, when they show a full-blown disease associated with kidney failure^11^. To evaluate the potential of CR in ameliorating disease phenotype, at the end of the CR experiment, 48-week-old CR Tg*^Umod^*^C147W^ mice were compared with 24- and 48-week-old AL Tg*^Umod^*^C147W^ and Tg*^Umod^*^wt^ mice (**Figure 5A**).

**Figure 5.**
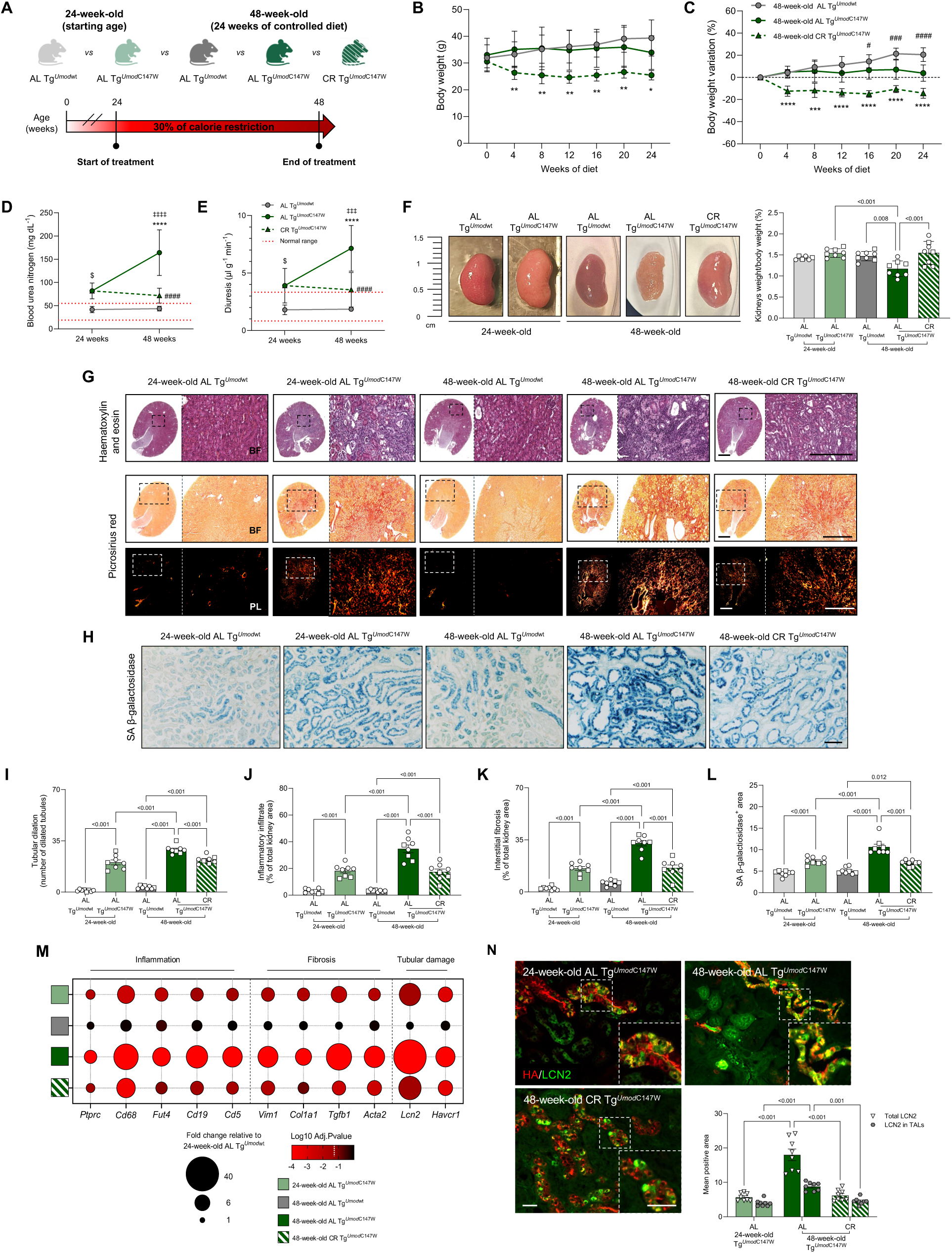
Effect of CR on ADTKD-*UMOD* progression at advanced disease stage. **A**) Schematic protocol of CR treatment at advanced disease stage. **B, C**) Body weight (**B**) and percentage of weight loss compared to the initial timepoint (**C**) of mice during the diet. ^#^*P*<0.05, ^###^*P*<0.001, ^####^*P*<0.0001 48-week-old AL Tg*^Umod^*^wt^ vs 48-week-old AL Tg*^Umod^*^C147W^; \**P*<0.05, \*\**P*<0.01, \*\*\**P*<0.001, \*\*\*\**P*<0.0001 48-week-old AL Tg*^Umod^*^C147W^ vs 48-week-old CR Tg*^Umod^*^C147W^. **D, E**) BUN (**D**) and diuresis (**E**) levels for the indicated experimental groups. Red dashed lines represent normal range values. ^$^*P*<0.05 24-week-old AL Tg*^Umod^*^wt^ vs 24-week-old AL Tg*^Umod^*^C147W^; \*\*\*\**P*<0.0001 48-week-old AL Tg*^Umod^*^wt^ vs 48-week-old AL Tg*^Umod^*^C147W^; ^‡‡‡^*P*<0.001, ^‡‡‡‡^*P*<0.0001 24-week-old AL Tg*^Umod^*^C147W^ vs 48-week-old AL Tg*^Umod^*^C147W^; ^####^*P*<0.0001 48-week-old AL Tg*^Umod^*^C147W^ vs 48-week-old CR Tg*^Umod^*^C147W^. **F**) Representative picture of kidneys from 24-week-old AL Tg*^Umod^*^wt^ and Tg*^Umod^*^C147W^, 48-week-old AL Tg*^Umod^*^wt^ and 48-week-old AL or CR Tg*^Umod^*^C147W^ mice and quantification of the kidney/body weight index. n=8 mice/group (5 female, 3 male). **G**) Representative histology images of kidney sections stained with haematoxylin and eosin or picrosirius red and viewed in bright field (BF, upper panels) or under polarized light (PL, bottom panels). Scale bars 2 mm and 600 μm. **H**) Representative images of senescence associated β-galactosidase (SA β-gal) staining of kidney sections. Scale bar 500 μm. **I, J, K, L**) Quantification of tubular dilation (**I**), inflammatory cell infiltrate (**J**), interstitial fibrosis (**K**) and SA β-gal positive areas (**L**). **M**) Categorical bubble plot showing the kidney expression levels of inflammation (*Ptprc*, *Cd68*, *Fut4*, *Cd19*, *Cd5*) fibrosis (*Vim1*, *Col1a1*, *Tgfb1*, *Acta2*) and tubular damage (*Lcn2*, *Havcr1*) related genes. Transcript levels were analysed by real-time RT-qPCR. Bubble size is proportional to fold change relative to 24-week-old AL Tg*^Umod^*^wt^ mice; bubble colour corresponds to Log_10_ of the adjusted P value compared to 24-week-old AL Tg*^Umod^*^wt^ mice (one-way ANOVA followed by Dunnett’s multiple comparisons post-hoc test). A Log_10_ adjusted *P* value ≤ −1.3 (white dashed line) was considered statistically significant. Data used to generate the categorical bubble plot are reported in *Supplemental Table 9*. **N**) Immunofluorescence staining for transgenic UMOD (HA, in red) and LCN2 (in green) in kidney sections. Histograms represent quantification of the LCN2 positive signal in total kidney area or in TAL tubules (HA^+^). n=8 mice group (5 females, 3 males). Data are reported as mean ± s.d. (in **B, C, D, E, F**) or ± s.e.m. (in **I, J, K, L, N**). Group comparisons were performed by using one-way ANOVA followed by Šídák’s correction (in **F, I, J, K, L**) or two-way ANOVA followed by Tukey’s correction (in **N**).

The prolonged CR protocol led to a relatively mild reduction of mice body weight (about 15%) (**Figure 5B, C**). We characterized kidney function through urine and plasma analysis. As expected, 24-week-old AL Tg*^Umod^*^C147W^ mice showed already compromised kidney function that significantly worsened with aging. Instead, such drop of kidney function is not observed in 48-week-old CR Tg*^Umod^*^C147W^ mice showing levels of plasma creatinine and BUN, urinary creatinine and diuresis comparable to 24-week-old mutant mice (**Figure 5D, E** and **Supplemental Table 7, 8**). The beneficial effect of the CR protocol was also evident when comparing kidney morphology at the end of treatment as the pale, shrunk appearance of kidneys of 48-week-old AL Tg*^Umod^*^C147W^ mice, resembling patient small-sized kidneys, was remarkably rescued (**Figure 5F**). In line with kidney function and morphology, we observed progressive worsening of disease parameters in 48-week-old AL Tg*^Umod^*^C147W^ mice compared with 24-week-old AL Tg*^Umod^*^C147W^ and Tg*^Umod^*^wt^ mice, that is essentially stopped by CR. This was observed for all investigated markers of inflammation, fibrosis and kidney damage, as assessed by histological (**Figure 5G, I-K**), immunofluorescence (**Supplemental Figure 4**) and gene expression analyses (**Figure 5M** and **Supplemental Table 9**). Since this experiment included aged mice, we also evaluated the kidney activity of senescence-associated β-galactosidase, recently reported as a predictor of renal outcome in patients with CKD of different origin.^47^ Interestingly, senescence was already increased in 24-week-old Tg*^Umod^*^C147W^ mice, relative to Tg*^Umod^*^wt^ mice. Also in this case, CR was effective in blocking further increase of senescence that is instead observed in 48-week-old AL Tg*^Umod^*^C147W^ mice (**Figure 5H, L**).

We also evaluated the effect of CR on preserving TAL integrity. In fact, compared to 24-week-old AL Tg*^Umod^*^C147W^ mice, 48-week-old AL Tg*^Umod^*^C147W^ mice show altered morphology of TAL segments and decreased expression the Na^+^-K^+^-2Cl^−^ co-transporter (NKCC2, marker of TAL cells) (**Supplemental Figure 5A-E**). We also observed significant increase of LCN2 in mutant uromodulin expressing cells (**Figure 5N** and **Supplemental Figure 5F**). All these parameters suggest progressive cell damage and possibly TAL cell loss in kidneys of 48-week-old AL Tg*^Umod^*^C147W^ mice, that is likely the reason why we did not observe further increase of mutant uromodulin aggregates compared to 24-week-old AL Tg*^Umod^*^C147W^ mice (**Supplemental Figure 5A**). Importantly, in 48-week-old CR Tg*^Umod^*^C147W^ mice we observed preserved TAL morphology (**Supplemental Figure 5C**), rescued expression levels of NKCC2 (**Supplemental Figure 5D, E**) and reduced LCN2 expression (**Figure 5N, Supplemental Figure 5F**), suggesting a role of CR in ameliorating progressive TAL damage in Tg*^Umod^*^C147W^ mice.

In sum, our data demonstrate that initiating CR at a later stage of the disease can still be effective in blocking ADTKD-*UMOD* progression.

### Unsupervised multivariate analysis of the overall therapeutic effect of CR

Finally, to integrate the values obtained for the characterization of the effect of CR on ADTKD-*UMOD* onset and progression, we performed unsupervised multivariate analysis. PCA followed by K-means clustering was conducted using 15 disease parameters evaluated in 8-week-old AL Tg*^Umod^*^wt^ and Tg*^Umod^*^C147W^ mice, and in 23-week-old AL or CR Tg*^Umod^*^wt^ and Tg*^Umod^*^C147W^ mice (**Figure 6A**). The heatmap showing the scaled value of all evaluated parameters in each sample is reported in **Supplemental Figure 6A**. PCA highlighted a very strong and statistically significant separation between all experimental groups (F-value 148.19). K-means analysis revealed separation of 8-week-old AL Tg*^Umod^*^C147W^ and 23-week-old AL Tg*^Umod^*^C147W^ groups into two distinct clusters (cluster 1 and 3, respectively), consistent with the progression of disease severity with age. Instead, all Tg*^Umod^*^wt^ groups formed a single cluster (cluster 4), clearly separated from the previous two. Notably, 23-week-old CR Tg*^Umod^*^C147W^ mice formed a fourth cluster (cluster 2), distinct from the other clusters composed by mutant mice, and more similar to wild-type animals (**Figure 6A** and **Supplemental Figure 6A**). This analysis substantiates the experimental evidence that, at early disease stages, CR blocks ADTKD-*UMOD* progression and reverts the already established kidney damage.

**Figure 6.**
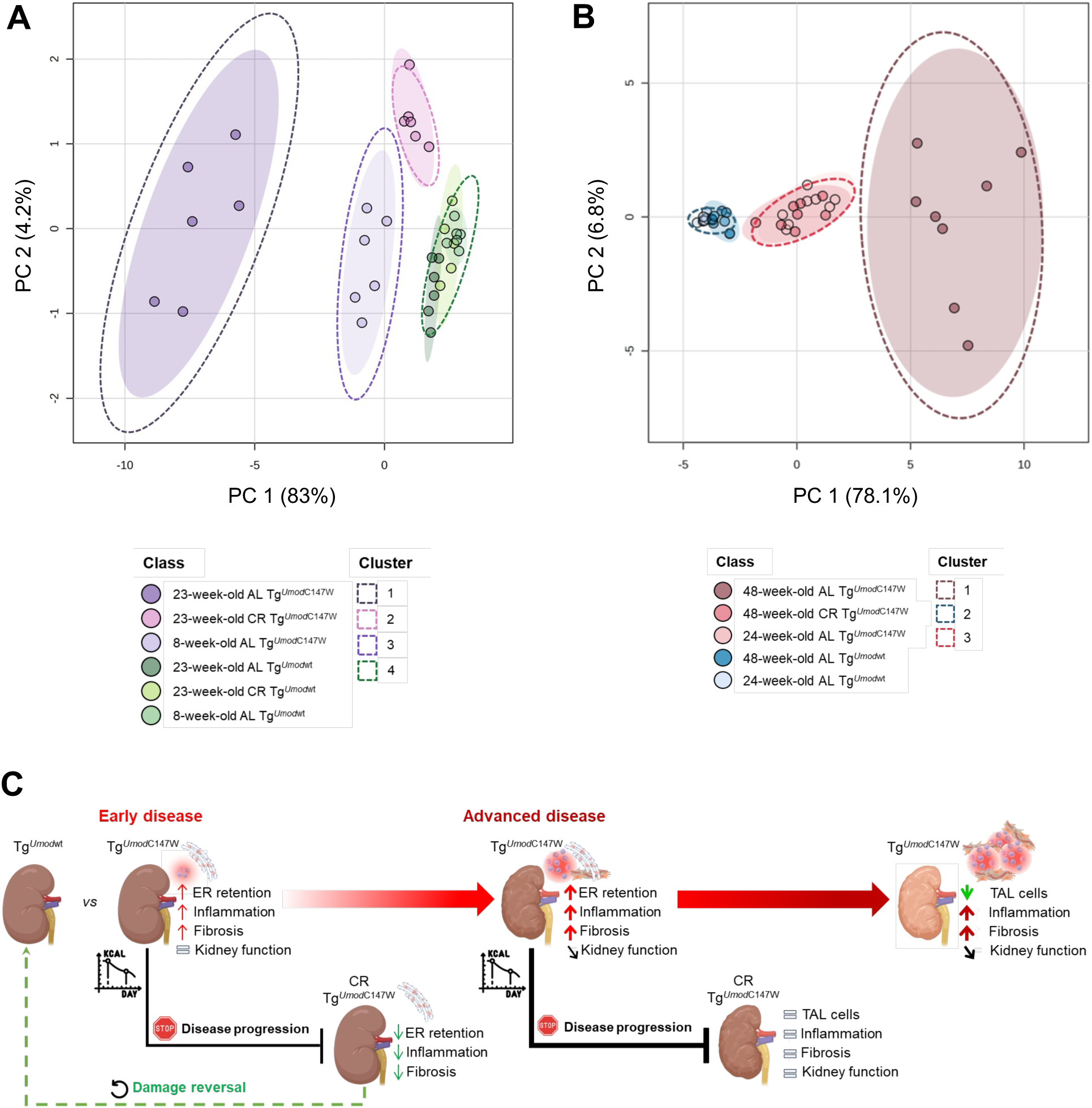
Summary of the effect of CR at early and advanced ADTKD-*UMOD* stages. **A**) PCA performed using 15 parameters measured in 8-week-old AL Tg*^Umod^*^wt^ or Tg*^Umod^*^C147W^ mice and 23-week-old AL or CR Tg*^Umod^*^wt^ and Tg*^Umod^*^C147W^ mice. A heatmap of the parameters used for the analysis is shown in *Supplemental* Figure 6A. F-value: 148.19; R-squared: 0.96; *P*<0.001 (based on 999 permutations) (PERMANOVA). K-means clustering of samples revealed 4 distinct clusters: cluster 1 (in black) is formed by 23-week-old AL Tg*^Umod^*^C147W^ mice, cluster 2 (in pink) is formed by 23-week-old CR Tg*^Umod^*^C147W^ mice, cluster 3 (in purple) is formed by 8-week-old AL Tg*^Umod^*^C147W^ mice while 8-week-old AL Tg*^Umod^*^wt^, 23-week-old AL Tg*^Umod^*^wt^ and 23-week-old CR Tg*^Umod^*^wt^ mice are all included in cluster 4 (in green). n=6 mice/group (all females). **B**) PCA performed using 21 parameters measured in 24-week-old AL Tg*^Umod^*^wt^ or Tg*^Umod^*^C147W^, 48-week-old AL Tg*^Umod^*^wt^ and 48-week-old AL or CR Tg*^Umod^*^C147W^ mice. A heatmap of the parameters used for the analysis is shown in *Supplemental* Figure 6B. F-value: 63.45; R-squared: 0.87; *P*<0.001 (based on 999 permutations) (PERMANOVA). K-means clustering of samples, based on the same disease parameters revealed 3 distinct clusters: cluster 1 (in brown) is formed by 48-week-old AL Tg*^Umod^*^C147W^ mice, cluster 2 (in blue) is formed by 24-week-old AL and 48-week-old AL Tg*^Umod^*^wt^ mice, cluster 3 (in red) is formed by 24-week-old AL Tg*^Umod^*^C147W^ and 48-week-old CR Tg*^Umod^*^C147W^ mice. n=8 mice/group. Each point represents an individual animal, color-coded according to its experimental group. Ellipses represent the 95% confidence interval area of each group. Dashed ellipses indicate 95% confidence intervals for each cluster centroid. **C**) Schematic representation summarizing the phenotypic effects induced by CR in Tg*^Umod^*^C147W^ mice at early and advanced disease stages.

A similar analysis was also performed to integrate the values of the 21 parameters tested in 24-week-old AL Tg*^Umod^*^wt^ and Tg*^Umod^*^C147W^ mice, and in 48-week-old AL Tg*^Umod^*^wt^ or AL and CR Tg*^Umod^*^wt^ and Tg*^Umod^*^C147W^ mice. PCA analysis demonstrates a strong and statistically significant difference between groups (F-value 63.45) (**Figure 6B**). Both the PCA results, and the K-means analysis revealed that wild-type mice cluster together, irrespectively of their age (24- and 48-week-old AL Tg*^Umod^*^wt^ mice, cluster 2) and are clearly separated from aged mutant mice fed AL (48-week-old AL Tg*^Umod^*^C147W^ mice, cluster 1). Instead, 48-week-old CR Tg*^Umod^*^C147W^ mice grouped together with 24-week-old AL Tg*^Umod^*^C147W^ mice (cluster 3) and are more similar to wild-type mice than their AL counterpart (**Figure 6B** and **Supplemental Figure 6B**). This reflects the experimental observations that CR prevents age-associated worsening of disease parameters.

Overall, these analyses provide additional and unsupervised evidence supporting the remarkable therapeutic efficacy of CR at both early and advanced ADTKD-*UMOD* stages (**Figure 6C**).

## Discussion

This study demonstrates the effectiveness of CR in reducing the intracellular load of mutant uromodulin and ameliorating kidney damage in ADTKD-*UMOD*. A 30% CR regimen for 15 or 24 weeks, starting at different stage of the disease^11,24^ blocks disease onset and progression in Tg*^Umod^*^C147W^ mice.

The beneficial effect of CR is likely downstream of CR-restored autophagy in TECs expressing mutant uromodulin, inducing degradation of ER-retained uromodulin and eventually preventing inflammation, fibrosis and kidney function decline. Our results have clear translational relevance by providing *in vivo* evidence of a possible therapeutic strategy for this rare, inherited cause of chronic kidney disease, for which specific therapies are still missing.^1,2^ Moreover, we provide the first *in vivo* evidence of the therapeutic value of a dietary intervention in managing an ER storage disease, potentially having implications beyond ADTKD.

The common primary effect of *UMOD* mutations on mutant uromodulin ER retention, as well as its downstream gain-of-toxic-function effect in disease pathogenesis are well established.^1^ Hence, interventions aimed at reducing mutant uromodulin intracellular load could be a valuable therapeutic option in the context of ADTKD-*UMOD*. Recently, a strategy aimed at rescuing mutant uromodulin trafficking, increasing its secretion, was shown to be effective in ameliorating kidney inflammation and fibrosis in an ADTKD-*UMOD* mouse model.^25^ However, the possible consequences of long-term secretion of mutant uromodulin have not been investigated. Here, we instead employed a strategy aimed at degrading mutant uromodulin through induction of autophagy prompted by CR.

Different studies describe CR as the most powerful non-genetic intervention to delay age-associated pathologies and extend lifespan in a large variety of species^26^, owing to its action on multiple molecular mechanisms, including enhancement of proteostasis, modulation of nutrient-sensing^29^ and autophagy induction. The effect of CR on mutant uromodulin degradation is demonstrated by remarkable reduction of uromodulin protein level, in absence of any effect on gene expression or urinary secretion. Consistent with previous results in other ADTKD-*UMOD* mouse models^14,19,22^, we reported the presence of P62 punctae in TALs of Tg*^Umod^*^C147W^ mice, together with increased total kidney levels of LC3-II/LC3-I ratio. This is likely downstream of mTOR activation, as demonstrated by increased levels of phospho-S6RP, suggesting defective autophagy in cells expressing mutant uromodulin. Upon CR, P62 aggregates and phospho-S6RP are strongly decreased in TALs, along with ER-retained mutant uromodulin. These results indicate that the beneficial role of CR is likely due to quenching of mTOR overactivation, a mechanism already described in mouse models of Autosomal Dominant Polycystic Kidney Disease (ADPKD) under mild to moderate (10-40%) CR.^48^ In Tg*^Umod^*^C147W^ mice, CR-mediated mTOR inhibition restores autophagy in TALs, likely leading to the clearance of mutant uromodulin. Our findings are consistent with previous *in vitro* and *in vivo* evidence. In cell models, autophagy stimulation using cell starvation or treatment with mTOR inhibitors (i.e., rapamycin or Torin-1 treatments), leads to intracellular removal of mutant uromodulin.^14,19^ In two different ADTKD-*UMOD* mouse models, autophagy induction via overexpression of MANF was shown to have beneficial effects on disease phenotype.^22^ Upon CR, clearance of mutant uromodulin aggregates is likely mediated by selective autophagy of the ER (ER-phagy)^49^ rather than bulk macro-autophagy. In kidneys of Tg*^Umod^*^C147W^ mice the ER-phagy receptors *Retreg1*, *Sec62* and *Tex264* were significantly downregulated, suggesting that ER-phagy is impaired in kidneys of mutant mice, failing to degrade ER-retained uromodulin aggregates. Upon CR, their transcript levels were recovered, likely reflecting a role of CR in restoring ER-phagy. *Retreg1* encodes for FAM134B, that plays a key role in both constitutive and starvation-induced ER turnover.^50^ Its expression is tightly controlled by the nutrient-sensitive Sestrin2-mTORC1-TFEB/TFE3 axis^44,51^ and, in response to ER stress, FAM134B mediates scavenging of ERAD-resistant misfolded clients through interaction with the ER chaperone calnexin (CNX).^52,53^ An N-terminal-truncated, alternative isoform of FAM134B (*Fam134b-2*) was shown to be transcriptionally enhanced to regulate ER-phagy under nutrient depletion.^45^ This is confirmed in our models, as *Fam134b-2,* but not *Fam134b-1* (encoding the full-length protein), was indeed up-regulated by CR. TEX264 was implicated in ER remodelling during nutrient stress^54^. SEC62 is a component of the protein translocon machinery that has been found to aid the autophagic degradation of the ER membranes during the recovery from ER stress, a process named recov-ER-phagy.^55^ Their induction in CR-treated mutant mice suggests a role in ER remodelling, possibly reflecting the resolution of ER stress following clearance of misfolded uromodulin aggregates. Given our previous finding that mutant uromodulin interacts with CNX^18^ and stemming from the effect of CR, further studies are warranted to dissect the relevance of FAM134B-CNX mediated ER-phagy and/or the role of SEC62 and TEX264 in the context of mutant uromodulin degradation.

CR was also effective in significantly ameliorating kidney inflammation. This anti-inflammatory effect of CR in the context of ADTKD-*UMOD* could be an indirect consequence of reduced TECs stress, following reduction of mutant uromodulin retention in the ER. This is supported by reduced LCN2 expression in TAL cells of CR-treated Tg*^Umod^*^C147W^ mice. However, a synergistic effect due to the direct anti-inflammatory role of CR is likely to play a role. Indeed, several studies demonstrated the potent capability of CR in modulating inflammation in different settings, ranging from inflammaging to inflammation of metabolic disorders and chronic diseases^27,56^, due to its direct impact on inflammatory cell metabolism.^35,36^ CR had a remarkable impact on kidney fibrosis. This is likely due to the effect on inflammation, in line with previous evidence highlighting the importance of inflammation in ADTKD-*UMOD* onset, and the potential therapeutic relevance of its targeting.^19,24^

The observation that in 23-week-old CR Tg*^Umod^*^C147W^ mice, inflammation and fibrosis markers are reduced to levels that are even lower that the ones observed at the start of dietary protocol (8 weeks), and comparable to wild-type mice, suggests that at early stages of disease CR is effective not only to block progression, but also to repair the already established kidney damage. Recently, fasting-mimicking diet, a dietary intervention that mimics the effects of fasting without requiring complete food restriction, was shown to restore normal nephron structure and function in a glomerulopathy model via the initiation of stem and nephrogenic pathways, thereby promoting podocyte regeneration.^57^ Whether the repair of kidney damage observed in our model is induced by similar pathways activated by CR remains to be tested. Kidney plasticity and ability to restore structural and functional changes have been described in a mouse model of ADPKD, showing reversal of disease phenotype following re-expression of Pkd genes. Recovery of kidney damage appeared complete when re-expression of Pkd occurred at early stage of disease, while it was only partial, with persisting signs of kidney scarring, if re-expression was induced at later time points.^58^ Similarly, our data show that at advanced disease stages (starting at 24 weeks of age and lasting till 48 weeks of age) CR is still effective in preventing worsening of inflammation, fibrosis and loss of kidney function, even if it does not revert the already established kidney phenotype. In 48-week-old AL Tg*^Umod^*^C147W^ mice we observed a marked reduction of NKCC2 expression with most TAL tubules showing dilation, flattened epithelia and increased LCN2 expression. This likely reflects progressive TAL cell damage and loss that we described to start at 24 weeks of age.^11^ In contrast, CR preserved the typical cuboidal morphology of TAL cells, maintained NKCC2 expression and reduced LCN2 levels. These findings suggest that CR effectively preserves the integrity of mutant uromodulin expressing cells and mitigates tubular damage at advanced stages of disease. In kidneys of 48-week-old Tg*^Umod^*^C147W^ mice we also reported increased activity of the SA β-gal, a well-established marker of cell senescence.^59^ Senescence has been described as a predictor of renal outcome in patients with CKD of diverse aetiology and a factor that exacerbates inflammation and fibrosis.^47,60^ In this context, CR has been reported as a powerful intervention to delay senescence and to improve changes associated with cellular senescence within the kidney, as tubular atrophy, epithelial to mesenchymal transition and interstitial fibrosis.^61,62^ In line with this, we observed a strong reduction of SA β-gal activity in kidneys of CR Tg*^Umod^*^C147W^ mice, suggesting that the pleiotropic action of CR might also include an effect on senescence. Further investigations will be needed to clarify the possible role of senescence in ADTKD-*UMOD* pathogenesis and its targeting as potential therapeutic intervention. Overall, considering the relatively late onset of kidney failure in ADTKD-*UMOD*, the finding that CR is effective in slowing disease progression, even when initiated at a later time point, could be relevant for patient management, as it might delay the need of replacement therapies.

Clinical trials investigating CR as an adjuvant therapy are currently ongoing in different disease settings, including multiple sclerosis^63^, ADPKD with overweight or obesity^64^ and type 2 diabetes.^65^ A possible limitation of CR use in clinical settings is the patients’ compliance to strictly adhere routinely to dietary restriction and the challenge, from the clinician’s point of view, to obtain a valid measure to evaluate CR adherence.^66^ Alternative strategies could envision the use of alternative dietary interventions, as time-restricted feeding, intermittent fasting or restriction of specific macronutrients^67,68^ or of pharmaceutical agents mimicking the effects of CR, collectively referred to as CR mimetics.^69^ These approaches could increase patient adherence to therapy, though their effectiveness in the context of ADTKD-*UMOD* needs to be explored. Additional studies are required to identify end points and biomarkers to transfer CR or similar autophagy-enhancing strategies (CRMs or alternative dietary interventions) to patients.

In conclusion, this work demonstrates that CR could be a valuable therapeutic intervention to treat ADTKD-*UMOD* and possibly other storage diseases characterised by pathological protein accumulation in the ER, inflammation and fibrosis.

## Supplemental Material

This article contains the following supplemental material:

### Supplemental Methods

#### Supplemental Figures

- Supplemental Figure 1. Effect of CR on uromodulin in male Tg*^Umod^*^C147W^ mice.
- Supplemental Figure 2. Effect of CR on renal inflammation and fibrosis in male Tg*^Umod^*^C147W^ mice.
- Supplemental Figure 3. Effect of CR on inflammation, fibrosis and tubular damage in Tg*^Umod^*^C147W^ mice at early disease stage.
- Supplemental Figure 4. Effect of CR on kidney inflammation and fibrosis in Tg*^Umod^*^C147W^ mice at advanced disease stage.
- Supplemental Figure 5. Effect of CR on TAL cell integrity in Tg*^Umod^*^C147W^ mice at advanced disease stage.
- Supplemental Figure 6. Heatmap of disease parameters tested to evaluate the effect of CR at early and advanced disease stage.

#### Supplemental Tables

- Supplemental Table 1. List and sequences of primers used for mouse genotyping.
- Supplemental Table 2. Data used to generate the categorical bubble plot in Figure 2.
- Supplemental Table 3. Kidney expression levels of genes encoding ER-phagy receptors.
- Supplemental Table 4. Blood urea nitrogen and serum creatinine levels in 8-week-old AL Tg*^Umod^*^wt^ and Tg*^Umod^*^C147W^ mice and 23-week-old AL or CR Tg*^Umod^*^wt^ and Tg*^Umod^*^C147W^ mice.
- Supplemental Table 5. Diuresis and urinary creatinine levels in 8-week-old AL Tg*^Umod^*^wt^ and Tg*^Umod^*^C147W^ mice and 23-week-old AL or CR Tg*^Umod^*^wt^ and Tg*^Umod^*^C147W^ mice.
- Supplemental Table 6. Data used to generate the categorical bubble plot in Figure 4.
- Supplemental Table 7. Blood urea nitrogen and serum creatinine levels in 24-week-old AL Tg*^Umod^*^wt^ and Tg*^Umod^*^C147W^, 48-week-old AL Tg*^Umod^*^wt^ and 48-week-old AL or CR Tg*^Umod^*^C147W^ mice.
- Supplemental Table 8. Diuresis and urinary creatinine levels in 24-week-old AL Tg*^Umod^*^wt^ and Tg*^Umod^*^C147W^, 48-week-old AL Tg*^Umod^*^wt^ and 48-week-old AL or CR Tg*^Umod^*^C147W^ mice.
- Supplemental Table 9. Data used to generate the categorical bubble plot in Figure 5.
- Supplemental Table 10. List and sequences of primers used for gene expression analysis by SYBR Green real-time RT-qPCR.
- Supplemental Table 11. List of antibodies used for Western blot and/or immunofluorescence analysis.

## Acknowledgments

We acknowledge the San Raffaele facilities of Animal Histopathology and in particular Amleto Fiocchi for acquisition of histology images and of Animal biochemistry, and particularly Micol Ravà for mouse serum analysis. We thank Dr. Francesco Consolato, Maurizio De Fusco, Prof. Giorgio Casari and Prof. Carmine Settembre for fruitful and critical discussion. This work is dedicated to the memory of Dr. Claudia Tammaro.

## Funding

This work was supported by the Italian Ministry of Health (RF-2021-12372538) and Fondazione Telethon (GMR23T2167) grants to L.R. M.G.C. was supported by fellowships from the Italian Ministry of University and Research, Fondazione Centro San Raffaele – Fondazione Fronzaroli and Vita-Salute San Raffaele University.

## Author contributions

Conceptualization and study design: MGC, LR. Methodology: MGC, BP, BC, CS. Experimentation and data acquisition: MGC, MM, BP, BC, CS. Funding acquisition: LR. Project administration: MGC, LR. Supervision: MGC, LR. Writing: MGC, CS, LR. The present work was performed by MGC in partial fulfilment of the requirements for obtaining the PhD degree at Vita-Salute San Raffaele University, Milan, Italy.

## Competing interests

All authors declare that they have no competing interests.

## Data and materials availability

All data associated with this study are available in the main text or the Supplementary Materials.

## Notes

### Competing Interest Statement

The authors have declared no competing interest.

