## Supplemental Material for "Calorie restriction leads to degradation of mutant uromodulin and ameliorates inflammation and fibrosis in *UMOD*-related kidney disease"

Cratere *et al.*

**Table of contents:**


1. **Supplemental Methods**
2. **Supplemental Figures**
   - Supplemental Figure 1. Effect of CR on uromodulin in male $Tg^{\mathit{Umod}\mathrm{C147W}}$ mice.
   - Supplemental Figure 2. Effect of CR on renal inflammation and fibrosis in male $Tg^{\mathit{Umod}\mathrm{C147W}}$ mice.
   - Supplemental Figure 3. Effect of CR on inflammation, fibrosis and tubular damage in $Tg^{\mathit{Umod}\mathrm{C147W}}$ mice at early disease stage.
   - Supplemental Figure 4. Effect of CR on kidney inflammation and fibrosis in $Tg^{\mathit{Umod}\mathrm{C147W}}$ mice at advanced disease stage.
   - Supplemental Figure 5. Effect of CR on TAL cell integrity in $Tg^{\mathit{Umod}\mathrm{C147W}}$ mice at advanced disease stage.
   - Supplemental Figure 6. Heatmap of disease parameters tested to evaluate the effect of CR at early and advanced disease stage.
3. **Supplemental Tables**
   - Supplemental Table 1. List and sequences of primers used for mouse genotyping.
   - Supplemental Table 2. Data used to generate the categorical bubble plot in Figure 2.
   - Supplemental Table 3. Kidney expression levels of genes encoding ER-phagy receptors.
   - Supplemental Table 4. Blood urea nitrogen and serum creatinine levels in 8-week-old AL $Tg^{\mathit{Umod}\mathrm{wt}}$ and $Tg^{\mathit{Umod}\mathrm{C147W}}$ mice and 23-week-old AL or CR $Tg^{\mathit{Umod}\mathrm{wt}}$ and $Tg^{\mathit{Umod}\mathrm{C147W}}$ mice.
   - Supplemental Table 5. Diuresis and urinary creatinine levels in 8-week-old AL $Tg^{\mathit{Umod}\mathrm{wt}}$ and $Tg^{\mathit{Umod}\mathrm{C147W}}$ mice and 23-week-old AL or CR $Tg^{\mathit{Umod}\mathrm{wt}}$ and $Tg^{\mathit{Umod}\mathrm{C147W}}$ mice.
   - Supplemental Table 6. Data used to generate the categorical bubble plot in Figure 4.
   - Supplemental Table 7. Blood urea nitrogen and serum creatinine levels in 24-week-old AL $Tg^{\mathit{Umod}\mathrm{wt}}$ and $Tg^{\mathit{Umod}\mathrm{C147W}}$, 48-week-old AL $Tg^{\mathit{Umod}\mathrm{wt}}$ and 48-week-old AL or CR $Tg^{\mathit{Umod}\mathrm{C147W}}$ mice.
   - Supplemental Table 8. Diuresis and urinary creatinine levels in 24-week-old AL $Tg^{\mathit{Umod}\mathrm{wt}}$ and $Tg^{\mathit{Umod}\mathrm{C147W}}$, 48-week-old AL $Tg^{\mathit{Umod}\mathrm{wt}}$ and 48-week-old AL or CR $Tg^{\mathit{Umod}\mathrm{C147W}}$ mice.
   - Supplemental Table 9. Data used to generate the categorical bubble plot in Figure 5.

- Supplemental Table 10. List and sequences of primers used for gene expression analysis by SYBR Green real-time RT-qPCR.
- Supplemental Table 11. List of antibodies used for Western blot and/or immunofluorescence analysis.

### 4. Supplemental References

### 1. Supplemental Methods

#### Study design

This study was designed to investigate the effect of CR in the context of ADTKD-*UMOD*. To this end, transgenic mice expressing wild-type and C147W mutant uromodulin ($Tg^{\mathit{Umod}\mathrm{wt}}$ and $Tg^{\mathit{Umod}\mathrm{C147W}}$) were subjected to a moderate 30% CR regimen for 15 or 24 weeks starting at different disease stages (8-week-old pre-symptomatic phase or 24-week-old post-symptomatic phase). We validated the effect of CR on mutant uromodulin biology, autophagy and ER-phagy induction. Kidney function, inflammation, fibrosis and tubular damage markers were tested by gene/protein expression analysis, immunofluorescence and histology to assess the effect of treatment on kidney phenotype. Histological scoring was performed independently by two operators. All animal experiments were reviewed and approved by the Institutional Animal Care and Use Committee of the San Raffaele Institute and were performed in accordance with the institutional guidelines (protocols number 1368 and 1526). Sample sizes (n=5-8 mice/group) were determined by statistical planning, which was part of the approval process for animal experiments. Animals from the same litter were randomly assigned to different experimental groups. All the sample numbers per experimental groups are reported in the figure legends. The investigators were not blinded for the data analysis. No samples were excluded from any experiments. Further methodological details are available below.

#### Urine and plasma collection and analysis

At the end of dietary interventions and after appropriate training, urine samples were collected for 16-hours during night period, using individual metabolic cages (Techniplast, Varese, IT). During the period in metabolic cages, mice were provided with free access to tap water and AL or rationed food, according to their experimental group. Once collected, urine samples were spun at 1000 x g for 10 minutes to remove food remnants. Diuresis was calculated as ratio between total volume of urine collected (µL), mouse body weight (g) and time of collection (min). Urinary creatinine was measured using the Creatinine (urinary) Colorimetric Assay kit (Cayman, Ann Arbor, MI, USA) following manufacturer's procedures. Prior to sacrifice, and after profound anaesthesia, up to 500 µL of blood were collected from mice by retro-orbital bleeding using heparinized capillary tubes (Hirschmann Laborgeräte, Eberstadt, DE). Blood samples were then centrifuged at 3000 x g for 10 minutes and plasma collected. Blood urea nitrogen and plasma creatinine levels were measured by the Core Facility of Animal Biochemistry (San Raffaele Scientific Institute, Milan), using the automatic biochemistry analyser iLab Aries (ANTISEI, Athens, EL).

**Mouse kidney collection and processing**

At the end of the experiments and after profound anaesthesia, mice were sacrificed by cervical dislocation followed by decapitation to collect the kidneys. The right kidney was decapsulated, cut in half and immediately stored at -80°C for subsequent RNA and protein extraction. The left kidney was decapsulated, divided in half and processed for immunofluorescence and histological analysis. For immunofluorescence, tissue was fixed for 16 hours with 4% paraformaldehyde solution, incubated for 16 hours in 30% sucrose solution, embedded in optimum cutting temperature embedding medium (OCT, Bio-Optica, Milan, IT) and snap-frozen in a mixture of isopropanol and dry ice. For histological analysis, kidney tissue was fixed in 4% formaldehyde solution, dehydrated in an increasing scale of ethanol (70%, 90%, 95% and 100%, Carlo Erba, Milan, IT), clarified in xylene (Bio-Optica) and embedded in paraffin (Bio-Optica).

**RNA extraction and real-time qPCR analysis**

Total RNA was extracted by homogenization of half kidney in PRImeZOL$^{TM}$ RNA isolation reagent (Canvax Reagents, Villadolid, ES) following manufacturer's protocol. RNA concentration was determined by using NanoDrop One spectrophotomer (Thermofisher, Waltham, MA, USA) and RNA quality was assessed on agarose gel. Reverse transcription of 1 µg of RNA was performed using iScript$^{TM}$ gDNA Clear cDNA Synthesis Kit (Bio-Rad, Hercules, CA, USA) according to the manufacturer's procedures. Expression of genes of interests was analysed by real-time qPCR analysis on CFX96TM Touch Real-time PCR (Bio-Rad) using the qPCR Core kit for SYBR® Green I No ROX (Eurogentec, Liège, BE). Specific primers (sequences provided in **Supplemental Table 10**) were designed by using Primer 3 software and their amplification efficiency was determined by dilution curve analysis. Expression levels of genes of interest were normalized to the one of *Hprt1*. The relative mRNA expression was calculated following the ΔΔCT method[1].

**Protein extraction and Western blot analysis**

For protein extraction, half kidneys were homogenized on ice using a tissue grinder, in a buffer containing 50 mM This-HCl pH 7.4, 150 mM NaCl, 1% Triton X-100, 0.5% SDS, 60 mM n-octyl-β-D-glucopyranoside (Sigma-Aldrich, St. Louis, MO, USA), 10 mM sodium fluoride, 1mM sodium orthovadanate, 1mM glycerol phosphate, 1:1000 Protease-Inhibitor Cocktail (Sigma-Aldrich). Kidney homogenates were incubated at 4°C for 60 min, under rotation and then centrifuged for 10 min at 17900 x g to separate the insoluble material. Protein concentration was assessed using Bradford protein assay (Biorad). Protein lysates or urine samples were separated by SDS-PAGE in reducing (using a loading buffer containing 100 mM dithiothreitol - DTT, Sigma-Aldrich - as reducing agent) or non-reducing condition and transferred on nitrocellulose membrane (GE-Heltcare, Chicago, IL, USA). After blocking, membranes were probed overnight with the indicated primary antibody (listed in **Supplemental Table 11**). Horseradish peroxidase-conjugated secondary antibodies (dilution 1:5000, GE-Heltcare, listed in **Supplemental Table 11**) were added and incubated for 90 minutes at room temperature. Protein bands were visualized with the Immobilon Western Chemiluminescent Horseradish Peroxidase Substrate kit (Millipore, Billerica, MA, USA) at Uvitec Alliance mini HD9 (Uvitec Imaging Systems, Cambridge, UK). Quantification of the optical band densities was performed using the gel analysis option of ImageJ software[2]. The optical density of proteins of interest was normalized on the one of Gapdh or on the Ponceau S staining of the same membranes.

**Immunofluorescence analysis**

Kidney sections (7 μm-thick, OCT-embedded) were permeabilized twice for 15 min at room temperature in PBS Tween-20 10% and then blocked for 1 hour at room temperature with 10% donkey serum in PBS with 0.1% Tween-20 or 0.1% Triton-X 100. The sections were incubated overnight at 4 °C with the indicated primary antibodies (listed in **Supplemental Table 11**) diluted in PBS with 1% donkey serum and 0.1% Tween-20. The proper fluorescent-conjugated secondary antibody (dilution 1:500 Alexa Fluor, Thermofisher, listed in **Supplemental Table 11**) was incubated for 90 min at room temperature. Nuclei were counterstained with diamidino-2-phenylindole (DAPI) and then the slides were mounted using FluorSave Reagent (Calbiochem, San Diego, CA, USA). All tissue sections were visualized under a DM 5000B fluorescence upright microscope (Leica DFC480 camera, Leica DFC Twain Software, 20x/0.75 lens; Leica Microsystems, Deerfield, IL, USA) and identical acquisition parameters were ensured for the visualization of the same antibody in different kidney sections. Quantification of the fluorescent positive area was performed by adjusting the threshold with a specific function of ImageJ software and then measured with analyse tools[2]. The kidney F4/80$^+$ area was evaluated by quantifying 16 representative pictures acquired in the cortical (8) and the medullary (8) area. For the evaluation of Vimentin$^+$ area, 15 representative field were quantified (10 cortical and 5 medullary). The colocalization of HA or UMOD with CLR and the signal levels of LCN2, phospho-S6RP or P62 in HA positive tubules were quantified by measuring the area of overlapping fluorescence signal with an ImageJ specific tool on 6 representative fields per animal.

**Histological analysis**

Kidney sections (7 μm-thick, paraffin-embedded) were used to perform histological analysis. Briefly, after de-waxing and hydration in decreasing scale of ethanol (100%, 95%, 90%, 70%, Carlo Erba), samples were stained with Hematoxylin and Eosin (Histo-Line Laboratories, Milan, IT) or with Picrosirius Red (Abcam, Cambridge, UK) following manufacturer's instructions. After staining, kidney sections were dehydrated, cleared, and mounted. Brightfield images of stained sections were digitally acquired using an Aperio ePathology digital scanner (Leica Biosystems, Nussloch, DE). Polarized light imaged of Picrosirius-red staining were acquired using an Axiophot microscope (Zeiss, Baden, Wurttemberg, DE) equipped with a Leica DFC450 C camera (Leica Microsystems). Tubular dilation was quantified on 6 field per each H&E-stained section (3 cortical and 3 medulla, 20x magnification), as previously described [3,4]. Briefly, a grid composed of dots distant one from another by 13.625 μm was overlapped on the images and tubule were counted as dilated when more than two dots were present in their lumen. Area of inflammatory cell infiltration was quantified on 10 fields per each H&E-stained kidney section (5 cortical and 5 medulla, 40x magnification) by selecting regions of interest (ROI) using the ImageJ dedicated tool. Semiquantitative analysis of interstitial fibrosis was performed by measuring the collagen-positive area in the entire Picrosirius Red-stained kidney section (3 sections per animal) viewed under polarized light. Threshold adjustments were applied using a specific ImageJ function. All images were analysed independently by two operators and the average value for each sample was calculated.

**Senescence associated β-galactosidase activity on cryopreserved kidney sections**

Kidney sections (7 μm-thick, OCT-embedded) were used to perform senescence associated β-galactosidase staining using Senescence β-Galactosidase Staining Kit (Cell Signalling, Danvers, MA, USA), following manufacturer's procedures. Briefly, kidney sections were fixed for 15 min at room temperature, washed twice in PBS and then incubated overnight at 37°C in β-Galactosidase Staining Solution at pH 6. After two washes in PBS, sections were mounted using 70% glycerol and visualized using an Axiophot microscope (Zeiss) equipped with a Leica DFC450 C camera (Leica Microsystems). For each sample, 10 pictures at 2.5x magnification were acquired. β-galactosidase positivity was quantified by using the ImageJ Threshold tool, after colour deconvolution using the ImageJ plugin Colour Deconvolution2 v2.1; vector FastRed FastBlue DAB. All images were analysed independently by two operators and the average value for each sample was calculated.

### 2. Supplemental Figures

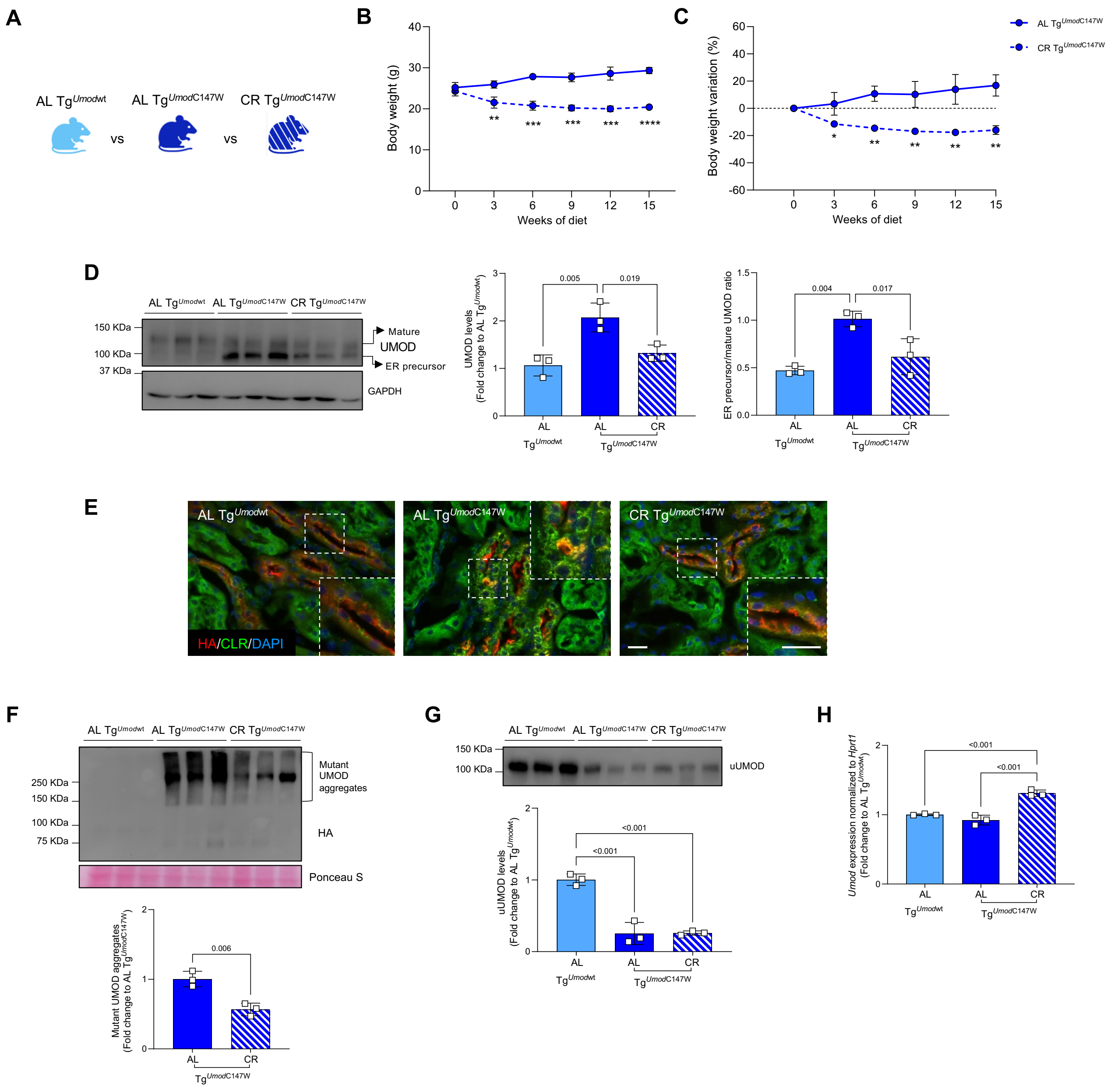

**Supplemental Figure 1. Effect of CR on uromodulin in male Tg$^{Umod\text{C147W}}$ mice. A**, **B**, **C**) Experimental groups (**A**), body weight (**B**) and body weight variation (expressed as percentage of body weight change compared to initial body weight) (**C**) of male Tg$^{Umod\text{C147W}}$ mice fed AL or CR during the diet. $^{*}P$<0.05, $^{**}P$<0.01, $^{***}P$<0.001, $^{****}P$<0.0001 (Unpaired t-test). **D**) Western blot analysis of UMOD in whole kidney homogenates. GAPDH is reported as a loading control. The histograms show the quantification of total UMOD and the ratio of ER precursor to mature UMOD isoforms. **E**) Immunofluorescence analysis for transgenic UMOD (HA, in red) and the ER marker CLR (in green). Nuclei are counterstained with DAPI (in blue). The magnified field is indicated by a dashed square on each picture. Scale bar 20 μm. **F**) Western blot analysis in non-reducing conditions and quantification of HMW mutant UMOD aggregates. **G**) Western blot analysis and quantification of UMOD urinary levels (uUMOD). Urine samples were normalized over total volume of a 16-hour urine collection in metabolic cage after appropriate training. **H**) Analysis (real-time RT-qPCR) of *Umod* expression in kidneys. Data are reported as fold change relative to AL $Tg^{\mathit{Umod}\mathrm{wt}}$ mice (in **D**, **G**, **H**) or to AL $Tg^{\mathit{Umod}\mathrm{C147W}}$ mice (in **F**). n = 3 mice/group. Data are reported as mean ± s.d. Group comparisons were performed by using one-way ANOVA followed by Tukey's correction (in **D**, **G**, **H**) or Unpaired t-test (in **F**).

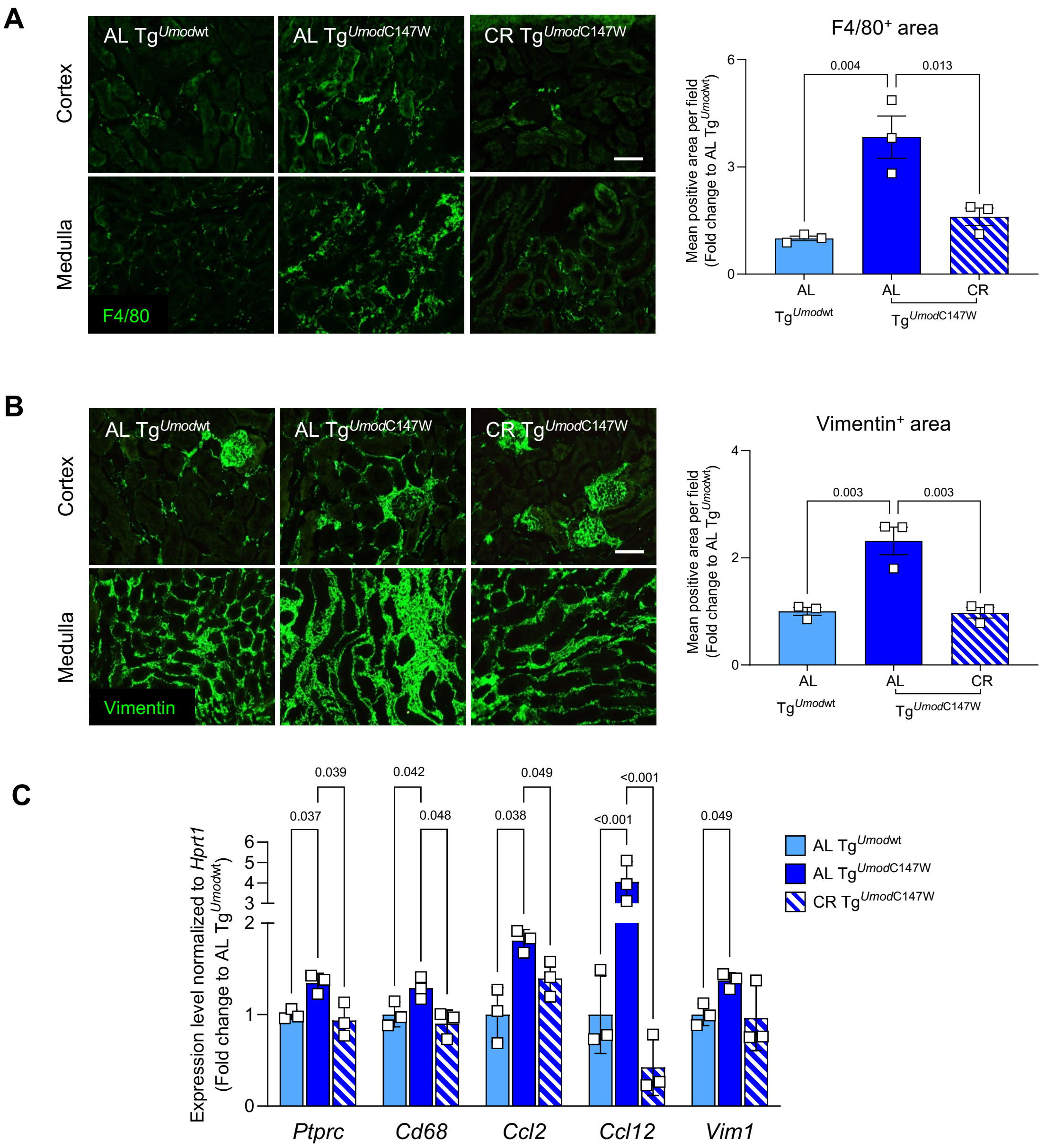


**Supplemental Figure 2. Effect of CR on renal inflammation and fibrosis in male Tg$^{UmodC147W}$ mice. A**, **B**) Immunofluorescence analysis and quantification of macrophage infiltration (F4/80) (**A**) and fibrosis (vimentin) (**B**) in cortical and medullary kidney portions. Scale bar 20 µm. **C**) Kidney expression level (real-time RT-qPCR) of genes encoding inflammatory cell markers (*Ptprc*, *Cd68*), pro-inflammatory chemokines (*Ccl2*, *Ccl12*) and fibrosis marker (*Vim1*). Data are reported as fold change relative to AL Tg$^{Umodwt}$ mice. n = 3 mice/group. Bars represent mean ± s.e.m. (in **A**, **B**) or ± s.d. (in **C**). Group comparisons were performed by using one-way ANOVA followed by Tukey's correction.

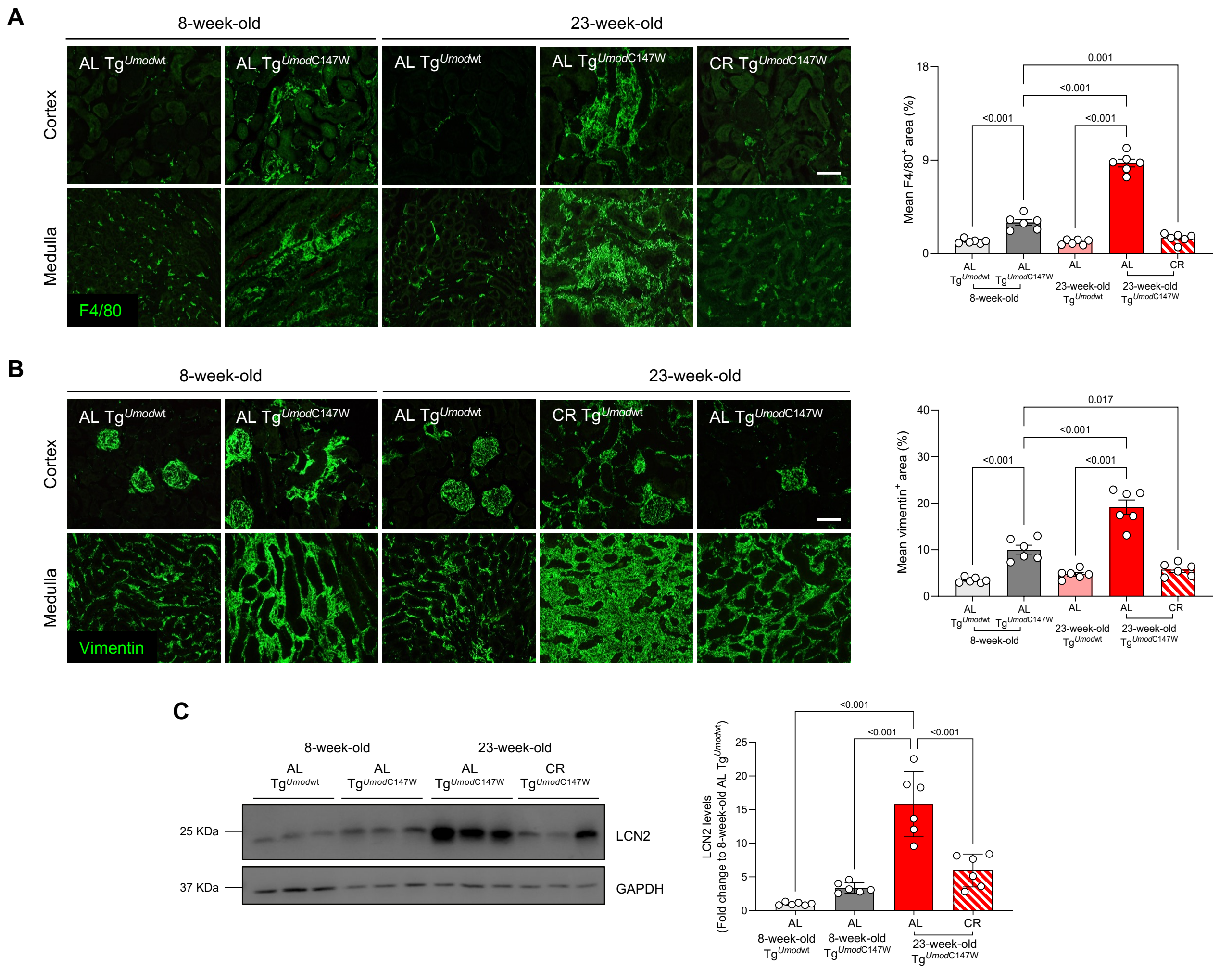


**Supplemental Figure 3. Effect of CR on inflammation, fibrosis and tubular damage in Tg$^{Umod C147W}$ mice at early disease stage. A**, **B**) Immunofluorescence analysis for F4/80 (**A**) and vimentin (**B**) in cortical and medullary kidney portions. Scale bar 20 μm. Histograms represent quantification of the kidney positive area for the indicated marker, expressed as percentage of total kidney area. n=6 animals/group. **C**) Western blot analysis and quantification of LCN2 levels in kidneys. GAPDH is reported as a loading control. Data are reported as mean ± s.e.m (in **A**, **B**) or ± s.d. (in **C**). Group comparisons were performed by using one-way ANOVA followed by Šídák's correction.

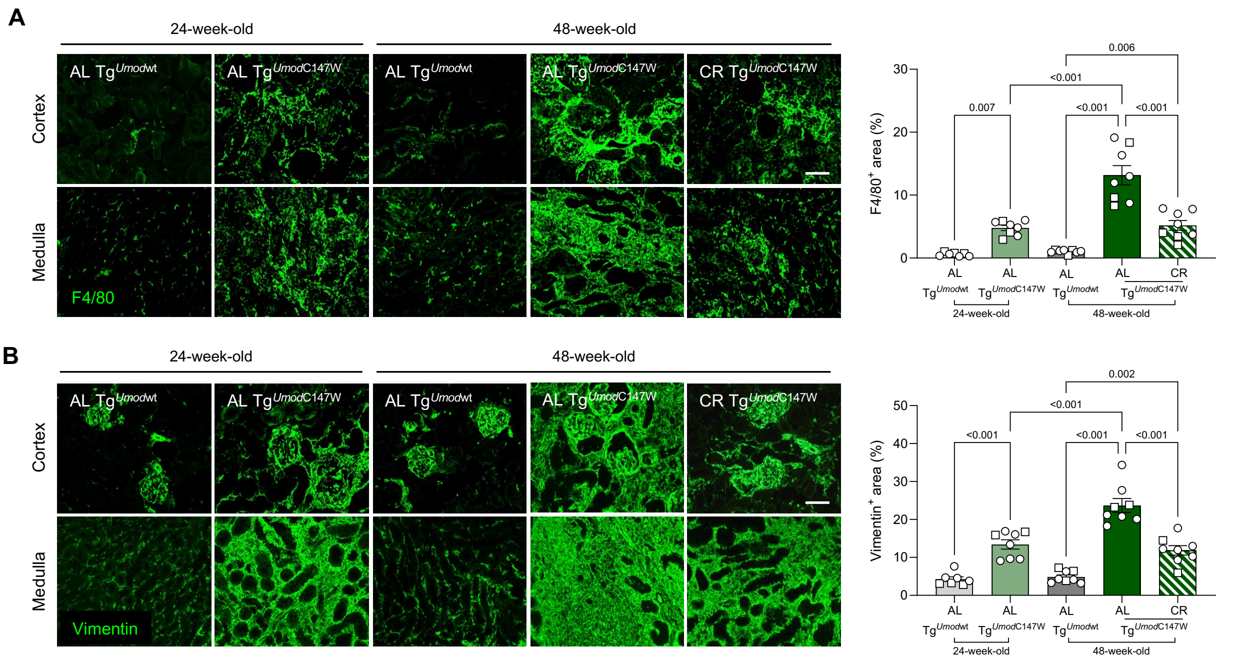


**Supplemental Figure 4. Effect of CR on kidney inflammation and fibrosis in Tg$^{\textit{Umod}\text{C147W}}$ mice at advanced disease stage. A, B**) Representative images of immunofluorescence analysis for F4/80 (**A**) and vimentin (**B**) in cortex and medullary kidney portions. Scale bar 20 µm. Histograms represent quantification of the kidney positive area for the indicated marker, expressed as percentage of total kidney area. n=8 animals/group. Data are reported as mean ± s.e.m. Group comparisons were performed by using one-way ANOVA followed by Šídák's correction.

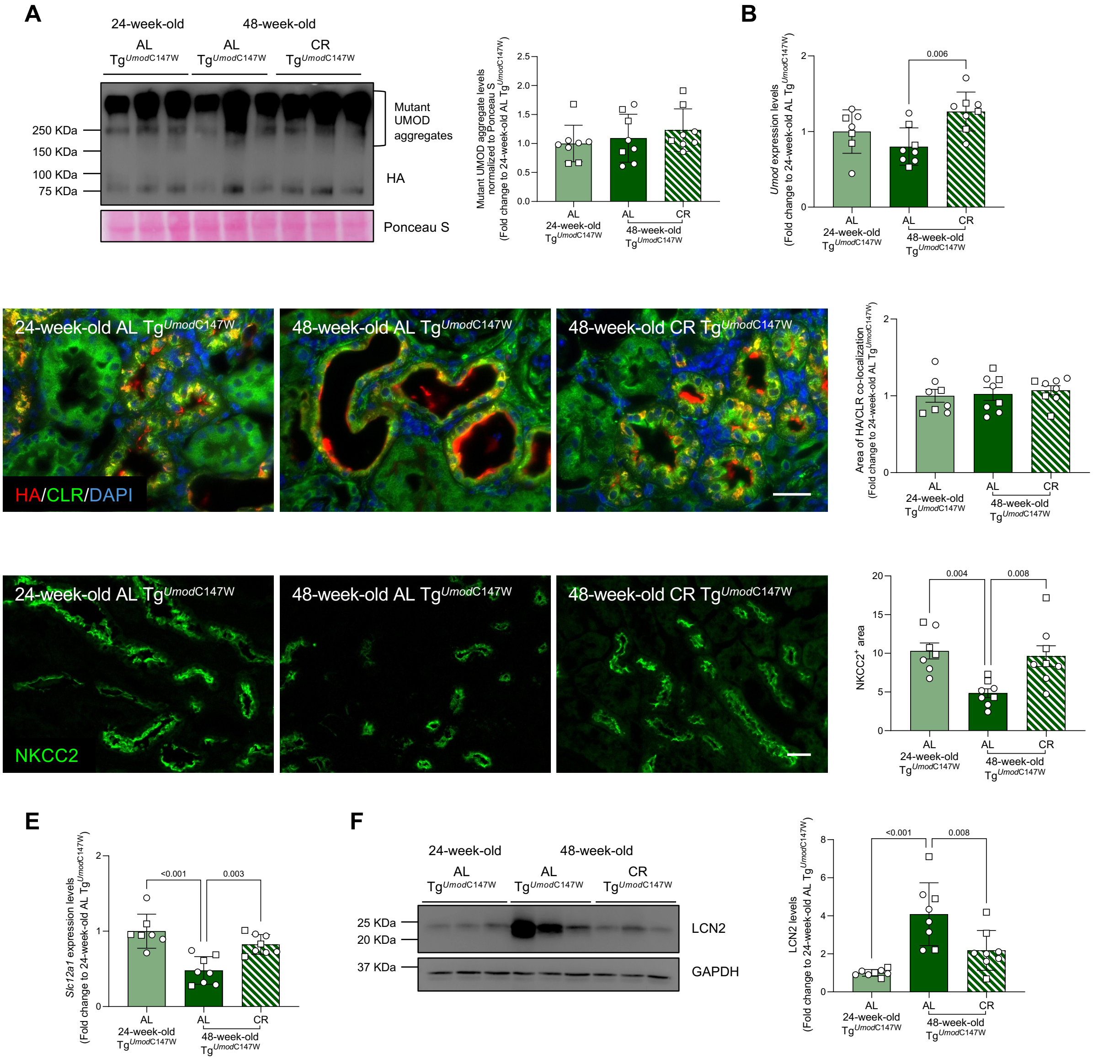


**Supplemental Figure 5. Effect of CR on TAL cell integrity in Tg$^{Umod\text{C147W}}$ mice at advanced disease stage. A**) Western blot in non-reducing condition showing transgenic UMOD (anti-HA antibody). Graph represents quantification of mutant UMOD aggregates. Ponceau S staining is shown as a loading control. **B**) Analysis by real-time RT-qPCR of *Umod* expression levels normalized over *Hprt1*. **C**) Immunofluorescence staining for transgenic UMOD (HA, in red) and CLR (in green) in kidneys of 24-week-old AL and 48-week-old AL or CR Tg$^{Umod\text{C147W}}$ mice. Nuclei are counterstained with DAPI (in blue). Scale bar 20 µm. Histograms represent quantification of the area of transgenic UMOD-CLR colocalization. **D**) Immunofluorescence staining for $Na^+$-$K^+$-$2Cl^-$ co-transporter (NKCC2) in kidney sections. Scale bar 20 µm. Histograms represent quantification of NKCC2 positive area. **E**) Analysis by real-time RT-qPCR of *Slc12a1* (encoding NKCC2) kidney expression levels. **F**) Western bot analysis and quantification of LCN2 levels in total kidney lysates. GAPDH is shown as a loading control. n=8 animals/group. Bars represent mean ± s.d. (in **A**, **B**, **E**, **F**) or mean ± s.e.m. (in **C**, **D**). In **A**, **B**, **C**, **E** and **F**, data are reported as fold change relative to 24-week-old AL $Tg^{\mathit{Umod}\,C147W}$ mice. Group comparisons were performed by using one-way ANOVA followed by Tukey's correction.

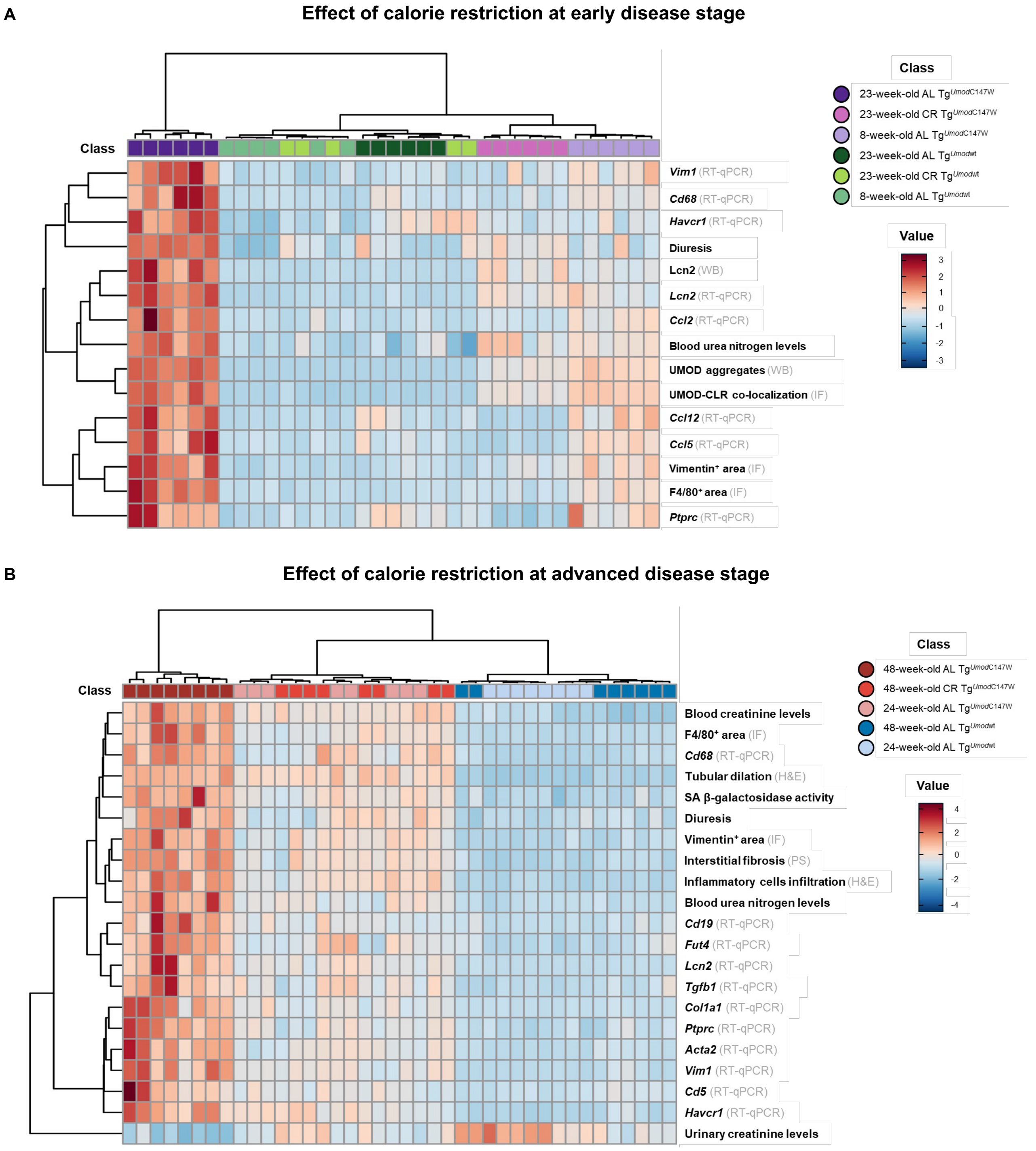


**Supplemental Figure 6. Heatmap of disease parameters tested to evaluate the effect of CR at early and advanced disease stage. A)** Heatmap showing the value of 15 disease parameters tested in the indicated experimental groups. These parameters were tested to evaluate the effect of CR on disease progression at early disease stage. The same data were used to perform PCA and K-means clustering analysis reported in *Figure*

*6A*. **B)** Heatmap showing the value of 21 disease parameters assessed to evaluate the effect of CR at advanced disease stage in the indicated experimental groups. The same parameters were used to perform PCA and K-means clustering analysis reported in *Figure 6B*. To ensure comparability between variables, data were normalized by auto-scaling transformation (mean-centered and divided by the s.d. of each variable), since disease features were measured according to different scales. RT-qPCR, real-time RT-qPCR; WB, Western blot; IF, immunofluorescence; H&E, haematoxylin and eosin staining; PS, picrosirius red staining.

### 3. Supplemental Tables

| Primer forward (5’ to 3’) | Primer reverse (5’ to 3’) |
|---|---|
| **Tg$^{Umod C147W}$ mouse line genotyping** | |
| GACAATGCTCAGGCTTCAGTG | GTCCAGAGGGTAAGAGCATTC |
| AAACTGCAGACCTAACCCATCCTTGCCTTTG | AAGTAGTCGCCTTCTGTGTTG |

**Supplemental Table 1.** List and sequences of primers used for mouse genotyping.

| Gene | Experimental group | Fold change relative to AL Tg$^{Umodwt}$ | Log$_{10}$ Adj. *P* value relative to AL Tg$^{Umodwt}$ |
|---|---|---|---|
| *Ptprc* | CR Tg$^{Umodwt}$ | 0.51 ± 0.12 | -1.75 |
| | AL Tg$^{UmodC147W}$ | 1.75 ± 0.46 | -3.52 |
| | CR Tg$^{UmodC147W}$ | 0.48 ± 0.12 | -2.04 |
| *Cd68* | CR Tg$^{Umodwt}$ | 0.64 ± 0.12 | -0.30 |
| | AL Tg$^{UmodC147W}$ | 3.35 ± 0.89 | -4.00 |
| | CR Tg$^{UmodC147W}$ | 0.79 ± 0.17 | -0.10 |
| *Fut4* | CR Tg$^{Umodwt}$ | 0.59 ± 0.20 | -0.66 |
| | AL Tg$^{UmodC147W}$ | 1.82 ± 0.36 | -4.00 |
| | CR Tg$^{UmodC147W}$ | 0.76 ± 0.40 | -0.97 |
| *Cd19* | CR Tg$^{Umodwt}$ | 0.61 ± 0.44 | -0.22 |
| | AL Tg$^{UmodC147W}$ | 2.48 ± 0.94 | -2.85 |
| | CR Tg$^{UmodC147W}$ | 0.57 ± 0.56 | -0.31 |
| *Cd5* | CR Tg$^{Umodwt}$ | 0.34 ± 0.17 | -0.92 |
| | AL Tg$^{UmodC147W}$ | 4.87 ± 1.54 | -3.15 |
| | CR Tg$^{UmodC147W}$ | 0.22 ± 0.09 | -0.35 |
| *Ccl2* | CR Tg$^{Umodwt}$ | 1.13 ± 0.08 | -0.14 |
| | AL Tg$^{UmodC147W}$ | 2.33 ± 0.41 | -4.00 |
| | CR Tg$^{UmodC147W}$ | 1.01 ± 0.23 | -0.004 |
| *Ccl12* | CR Tg$^{Umodwt}$ | 0.23 ± 0.18 | -0.23 |
| | AL Tg$^{UmodC147W}$ | 8.99 ± 2.18 | -4.00 |
| | CR Tg$^{UmodC147W}$ | 0.75 ± 0.57 | -0.01 |
| *Ccl5* | CR Tg$^{Umodwt}$ | 0.18 ± 0.11 | -2.89 |
| | AL Tg$^{UmodC147W}$ | 1.95 ± 0.55 | -3.70 |
| | CR Tg$^{UmodC147W}$ | 0.20 ± 0.10 | -3.00 |
| *Vim1* | CR Tg$^{Umodwt}$ | 0.84 ± 0.29 | -0.04 |
| | AL Tg$^{UmodC147W}$ | 2.64 ± 0.82 | -4.00 |
| | CR Tg$^{UmodC147W}$ | 0.77 ± 0.22 | -0.12 |
| *Tgfb1* | CR Tg$^{Umodwt}$ | 0.77 ± 0.31 | -0.33 |
| | AL Tg$^{UmodC147W}$ | 1.84 ± 0.44 | -3.40 |
| | CR Tg$^{UmodC147W}$ | 0.54 ± 0.22 | -1.32 |
| *Col1a1* | CR Tg$^{Umodwt}$ | 0.49 ± 0.30 | -0.14 |
| | AL Tg$^{UmodC147W}$ | 5.40 ± 1.84 | -4.00 |
| | CR Tg$^{UmodC147W}$ | 0.35 ± 0.15 | -0.27 |
| *Acta2* | CR Tg$^{Umodwt}$ | 1.21 ± 0.27 | -0.17 |
| | AL Tg$^{UmodC147W}$ | 2.10 ± 0.44 | -4.00 |
| | CR Tg$^{UmodC147W}$ | 0.48 ± 0.12 | -1.24 |
| *Lcn2* | CR Tg$^{Umodwt}$ | 0.71 ± 0.17 | -0.00004 |
| | AL Tg$^{UmodC147W}$ | 28 ± 7.43 | -4.00 |
| | CR Tg$^{UmodC147W}$ | 7.11 ± 6.09 | -0.98 |
| *Havcr1* | CR Tg$^{Umodwt}$ | 0.97 ± 0.47 | -0.0002 |
| | AL Tg$^{UmodC147W}$ | 3.46 ± 0.99 | -4.00 |
| | CR Tg$^{UmodC147W}$ | 0.86 ± 0.52 | -0.02 |

**Supplemental Table 2. Data used to generate the categorical bubble plot in Figure 2.** Fold change (mean ± s.d.) and $Log_{10}$ of the adjusted *P* value (one-way ANOVA followed by Dunnett's test) relative to AL $Tg^{\mathit{Umod}\mathrm{wt}}$ of the kidney expression level of the specified genes in the indicated experimental groups. A value ≤ -1.3 (corresponding to a *P* value ≤0.05) was considered statistically significant.

| Gene | Fold change relative to AL $Tg^{Umodwt}$ | | | *P* value | | | | |
|---|---|---|---|---|---|---|---|---|
| | CR $Tg^{Umodwt}$ | AL $Tg^{UmodC147W}$ | CR $Tg^{UmodC147W}$ | AL $Tg^{Umodwt}$ vs CR $Tg^{Umodwt}$ | AL $Tg^{Umodwt}$ vs AL $Tg^{UmodC147W}$ | AL $Tg^{Umodwt}$ vs CR $Tg^{UmodC147W}$ | AL $Tg^{UmodC147W}$ vs CR $Tg^{UmodC147W}$ | CR $Tg^{Umodwt}$ vs CR $Tg^{UmodC147W}$ |
| *Retreg1* | 1.48 ± 0.27 | 0.48 ± 0.14 | 1.04 ± 0.34 | $P<0.05$ | $P<0.01$ | ns | $P<0.05$ | $P<0.01$ |
| *Retreg2* | 1.04 ± 0.27 | 1.01 ± 0.20 | 0.90 ± 0.14 | ns | ns | ns | ns | ns |
| *Retreg3* | 0.99 ± 0.19 | 0.90 ± 0.10 | 0.98 ± 0.06 | ns | ns | ns | ns | ns |
| *Atl3* | 1.12 ± 0.35 | 1.15 ± 0.08 | 1.10 ± 0.31 | ns | ns | ns | ns | ns |
| *Rtn3* | 1.06 ± 0.21 | 1.14 ± 0.17 | 1.24 ± 0.20 | ns | ns | ns | ns | ns |
| *Sec62* | 1.02 ± 0.15 | 0.68 ± 0.18 | 1.10 ± 0.20 | ns | $P<0.05$ | ns | $P<0.01$ | ns |
| *Tex264* | 1.21 ± 0.12 | 0.83 ± 0.09 | 1.20 ± 0.11 | $P<0.01$ | $P<0.05$ | $P<0.01$ | $P<0.0001$ | ns |
| *Ccpg1* | 1.12 ± 0.41 | 0.58 ± 0.15 | 0.57 ± 0.10 | ns | $P<0.05$ | $P<0.05$ | ns | $P<0.01$ |

**Supplemental Table 3. Kidney expression levels of genes encoding ER-phagy receptors.** The table summarizes the fold change (mean ± s.d.) relative to AL $Tg^{Umodwt}$ and the *P* value for the specified comparisons (one-way ANOVA followed by Šídák's correction) of the kidney expression level for the indicated genes in CR $Tg^{Umodwt}$ and AL or CR $Tg^{UmodC147W}$ mice. A *P* value <0.05 is considered statistically significant. Mean expression levels of the analysed genes for each experimental group are also reported in *Figure 3G.*

| Blood urea nitrogen (mg dl $^{-1}$) | |
|---|---|
| Group | Mean ± s.d. |
| 8-week-old AL Tg$^{Umod\text{wt}}$ | 35.00 ± 2.45 |
| 8-week-old AL Tg$^{Umod\text{C147W}}$ | 47.17 ± 2.23 |
| 23-week-old AL Tg$^{Umod\text{wt}}$ | 36.00 ± 6.81 |
| 23-week-old CR Tg$^{Umod\text{wt}}$ | 31.60 ± 8.53 |
| 23-week-old AL Tg$^{Umod\text{C147W}}$ | 70.71 ± 6.24 |
| 23-week-old CR Tg$^{Umod\text{C147W}}$ | 48.83 ± 9.15 |
| Normal range values[5] | 19 – 55 |
| *P* value | |
| 8-week-old AL Tg$^{Umod\text{wt}}$ vs 8-week-old AL Tg$^{Umod\text{C147W}}$ | $P$<0.05 |
| 8-week-old AL Tg$^{Umod\text{wt}}$ vs 23-week-old AL Tg$^{Umod\text{wt}}$ | ns |
| 8-week-old AL Tg$^{Umod\text{C147W}}$ vs 23-week-old AL Tg$^{Umod\text{C147W}}$ | $P$<0.0001 |
| 23-week-old AL Tg$^{Umod\text{wt}}$ vs 23-week-old CR Tg$^{Umod\text{wt}}$ | ns |
| 23-week-old AL Tg$^{Umod\text{wt}}$ vs 23-week-old AL Tg$^{Umod\text{C147W}}$ | $P$<0.0001 |
| 23-week-old AL Tg$^{Umod\text{wt}}$ vs 23-week-old CR Tg$^{Umod\text{C147W}}$ | $P$<0.05 |
| 23-week-old AL Tg$^{Umod\text{C147W}}$ vs 23-week-old CR Tg$^{Umod\text{C147W}}$ | $P$<0.0001 |
| 8-week-old AL Tg$^{Umod\text{C147W}}$ vs 23-week-old CR Tg$^{Umod\text{C147W}}$ | ns |
| **Serum creatinine (mg dl $^{-1}$)** | |
| Group | Mean ± s.d. |
| 8-week-old AL Tg$^{Umod\text{wt}}$ | N/A |
| 8-week-old AL Tg$^{Umod\text{C147W}}$ | N/A |
| 23-week-old AL Tg$^{Umod\text{wt}}$ | 0.28 ± 0.01 |
| 23-week-old CR Tg$^{Umod\text{wt}}$ | 0.31 ± 0.07 |
| 23-week-old AL Tg$^{Umod\text{C147W}}$ | 0.36 ± 0.03 |
| 23-week-old CR Tg$^{Umod\text{C147W}}$ | 0.28 ± 0.01 |
| Normal range values[6] | 0.31 – 0.47 |
| *P* value | |
| 23-week-old AL Tg$^{Umod\text{wt}}$ vs 23-week-old AL Tg$^{Umod\text{C147W}}$ | $P$<0.05 |
| 23-week-old AL Tg$^{Umod\text{wt}}$ vs 23-week-old CR Tg$^{Umod\text{wt}}$ | ns |
| 23-week-old AL Tg$^{Umod\text{wt}}$ vs 23-week-old CR Tg$^{Umod\text{C147W}}$ | ns |
| 23-week-old AL Tg$^{Umod\text{C147W}}$ vs 23-week-old CR Tg$^{Umod\text{C147W}}$ | $P$<0.05 |

**Supplemental Table 4. Blood urea nitrogen and serum creatinine levels in 8-week-old AL Tg$^{Umod\text{wt}}$ and Tg$^{Umod\text{C147W}}$ mice and 23-week-old AL or CR Tg$^{Umod\text{wt}}$ and Tg$^{Umod\text{C147W}}$ mice.** Data are mean ± s.d. n = 5-6 mice/group. The *P* value for the specified comparisons (one-way ANOVA followed Šídák's correction) is reported. N/A, not assessed; ns, not significant.

| Diuresis (μl $gBW^{-1}$ $h^{-1}$) | |
|---|---|
| Group | Mean ± s.d. |
| 8-week-old AL $Tg^{Umod wt}$ | 1.74 ± 0.42 |
| 8-week-old AL $Tg^{Umod C147W}$ | 2.34 ± 0.60 |
| 23-week-old AL $Tg^{Umod wt}$ | 2.22 ± 0.60 |
| 23-week-old CR $Tg^{Umod wt}$ | 2.28 ± 0.54 |
| 23-week-old AL $Tg^{Umod C147W}$ | 4.74 ± 0.18 |
| 23-week-old CR $Tg^{Umod C147W}$ | 2.82 ± 0.30 |
| Normal range values[7] | 0.83 – 3.33 |
| *P* value | |
| 8-week-old AL $Tg^{Umod wt}$ vs 8-week-old AL $Tg^{Umod C147W}$ | ns |
| 8-week-old AL $Tg^{Umod wt}$ vs 23-week-old AL $Tg^{Umod wt}$ | ns |
| 8-week-old AL $Tg^{Umod C147W}$ vs 23-week-old AL $Tg^{Umod C147W}$ | $P<0.0001$ |
| 23-week-old AL $Tg^{Umod wt}$ vs 23-week-old CR $Tg^{Umod wt}$ | ns |
| 23-week-old AL $Tg^{Umod wt}$ vs 23-week-old AL $Tg^{Umod C147W}$ | $P<0.0001$ |
| 23-week-old AL $Tg^{Umod wt}$ vs 23-week-old CR $Tg^{Umod C147W}$ | ns |
| 23-week-old AL $Tg^{Umod C147W}$ vs 23-week-old CR $Tg^{Umod C147W}$ | $P<0.0001$ |
| 8-week-old AL $Tg^{Umod C147W}$ vs 23-week-old CR $Tg^{Umod C147W}$ | ns |
| **Urinary creatinine (μmol $gBW^{-1}$ $24h^{-1}$)** | |
| Group | Mean ± s.d. |
| 8-week-old AL $Tg^{Umod wt}$ | N/A |
| 8-week-old AL $Tg^{Umod C147W}$ | N/A |
| 23-week-old AL $Tg^{Umod wt}$ | 0.18 ± 0.04 |
| 23-week-old CR $Tg^{Umod wt}$ | 0.24 ± 0.07 |
| 23-week-old AL $Tg^{Umod C147W}$ | 0.09 ± 0.01 |
| 23-week-old CR $Tg^{Umod C147W}$ | 0.18 ± 0.03 |
| Normal range values[5] | 0.11 – 0.25 |
| *P* value | |
| 23-week-old AL $Tg^{Umod wt}$ vs 23-week-old AL $Tg^{Umod C147W}$ | $P<0.01$ |
| 23-week-old AL $Tg^{Umod wt}$ vs 23-week-old CR $Tg^{Umod wt}$ | ns |
| 23-week-old AL $Tg^{Umod wt}$ vs 23-week-old CR $Tg^{Umod C147W}$ | ns |
| 23-week-old AL $Tg^{Umod C147W}$ vs 23-week-old CR $Tg^{Umod C147W}$ | $P<0.01$ |

**Supplemental Table 5. Diuresis and urinary creatinine levels in 8-week-old AL $Tg^{Umod wt}$ and $Tg^{Umod C147W}$ mice and 23-week-old AL or CR $Tg^{Umod wt}$ and $Tg^{Umod C147W}$ mice.** Data are mean ± s.d. n=5-6 mice/group. The *P* value for the specified comparisons (one-way ANOVA followed Šídák's correction) is reported. N/A, not assessed; ns, not significant.

| Gene | Experimental group | Fold change relative to 8-week-old AL Tg$^{Umod wt}$ | $Log_{10}$ (adj *P* value) relative to 8-week-old AL Tg$^{Umod wt}$ |
|---|---|---|---|
| *Ptprc* | 8-week-old AL Tg$^{Umod C147W}$ | 3.18 ± 1.39 | -1.87 |
| | 23-week-old AL Tg$^{Umod C147W}$ | 5.28 ± 1.86 | -4.00 |
| | 23-week-old CR Tg$^{Umod C147W}$ | 0.81 ± 0.47 | -0.0066 |
| *Cd68* | 8-week-old AL Tg$^{Umod C147W}$ | 1.94 ± 0.26 | -2.43 |
| | 23-week-old AL Tg$^{Umod C147W}$ | 3.57 ± 0.81 | -4.00 |
| | 23-week-old CR Tg$^{Umod C147W}$ | 0.84 ± 0.14 | -0.065 |
| *Ccl2* | 8-week-old AL Tg$^{Umod C147W}$ | 2.26 ± 0.53 | -0.97 |
| | 23-week-old AL Tg$^{Umod C147W}$ | 6.07 ± 1.89 | -4.00 |
| | 23-week-old CR Tg$^{Umod C147W}$ | 1.05 ± 0.21 | -0.00017 |
| *Ccl5* | 8-week-old AL Tg$^{Umod C147W}$ | 2.34 ± 0.34 | -1.93 |
| | 23-week-old AL Tg$^{Umod C147W}$ | 4.84 ± 1.48 | -4.00 |
| | 23-week-old CR Tg$^{Umod C147W}$ | 0.61 ± 0.18 | -0.20 |
| *Ccl12* | 8-week-old AL Tg$^{Umod C147W}$ | 7.19 ± 3.15 | -3.04 |
| | 23-week-old AL Tg$^{Umod C147W}$ | 14.97 ± 3.82 | -4.00 |
| | 23-week-old CR Tg$^{Umod C147W}$ | 0.48 ± 0.33 | -0.014 |
| *Vim1* | 8-week-old AL Tg$^{Umod C147W}$ | 2.36 ± 0.77 | -1.61 |
| | 23-week-old AL Tg$^{Umod C147W}$ | 5.09 ± 1.15 | -4.00 |
| | 23-week-old CR Tg$^{Umod C147W}$ | 9.45 ± 0.78 | -0.15 |
| *Lcn2* | 8-week-old AL Tg$^{Umod C147W}$ | 31.59 ± 4.61 | -2.18 |
| | 23-week-old AL Tg$^{Umod C147W}$ | 10.47 ± 4.67 | -4.00 |
| | 23-week-old CR Tg$^{Umod C147W}$ | 2.81 ± 2.84 | -4.00 |
| *Havcr1* | 8-week-old AL Tg$^{Umod C147W}$ | 8.18 ± 0.87 | -2.03 |
| | 23-week-old AL Tg$^{Umod C147W}$ | 2.65 ± 1.55 | -4.00 |
| | 23-week-old CR Tg$^{Umod C147W}$ | 9.45 ± 0.49 | -1.74 |

**Supplemental Table 6. Data used to generate the categorical bubble plot in Figure 4.** Fold change (mean ± s.d.) and $Log_{10}$ of the adjusted *P* value (one-way ANOVA followed by Dunnett's correction) relative to 8-week-old AL Tg$^{Umod wt}$ of the kidney expression level of the indicated genes in 8-week-old AL Tg$^{Umod C147W}$ and AL or CR 23-week-old Tg$^{Umod C147W}$ mice. A value $\leq$-1.3 (corresponding to a *P* value $\leq$0.05) was considered statistically significant.

| Blood urea nitrogen (mg dl$^{-1}$) | |
|---|---|
| Group | Mean ± s.d. |
| 24-week-old AL Tg$^{Umod\text{wt}}$ | 41.62 ± 6.80 |
| 24-week-old AL Tg$^{Umod\text{C147W}}$ | 81.75 ± 17.06 |
| 48-week-old AL Tg$^{Umod\text{wt}}$ | 43.63 ± 4.92 |
| 48-week-old AL Tg$^{Umod\text{C147W}}$ | 164.13 ± 49.24 |
| 48-week-old CR Tg$^{Umod\text{C147W}}$ | 71.75 ± 15.93 |
| Normal range values[5] | 19 – 55 |
| *P* value | |
| 24-week-old AL Tg$^{Umod\text{wt}}$ vs 24-week-old AL Tg$^{Umod\text{C147W}}$ | *P*<0.05 |
| 24-week-old AL Tg$^{Umod\text{wt}}$ vs 48-week-old AL Tg$^{Umod\text{wt}}$ | ns |
| 24-week-old AL Tg$^{Umod\text{C147W}}$ vs 48-week-old AL Tg$^{Umod\text{C147W}}$ | *P*<0.0001 |
| 48-week-old AL Tg$^{Umod\text{wt}}$ vs 48-week-old AL Tg$^{Umod\text{C147W}}$ | *P*<0.0001 |
| 48-week-old AL Tg$^{Umod\text{wt}}$ vs 48-week-old CR Tg$^{Umod\text{C147W}}$ | ns |
| 48-week-old AL Tg$^{Umod\text{C147W}}$ vs 48-week-old CR Tg$^{Umod\text{C147W}}$ | *P*<0.0001 |
| 24-week-old AL Tg$^{Umod\text{C147W}}$ vs 48-week-old CR Tg$^{Umod\text{C147W}}$ | ns |
| **Serum creatinine (mg dl$^{-1}$)** | |
| Group | Mean ± s.d. |
| 24-week-old AL Tg$^{Umod\text{wt}}$ | 0.29 ± 0.03 |
| 24-week-old AL Tg$^{Umod\text{C147W}}$ | 0.39 ± 0.04 |
| 48-week-old AL Tg$^{Umod\text{wt}}$ | 0.28 ± 0.02 |
| 48-week-old AL Tg$^{Umod\text{C147W}}$ | 0.55 ± 0.10 |
| 48-week-old CR Tg$^{Umod\text{C147W}}$ | 0.38 ± 0.05 |
| Normal range values[6] | 0.31 – 0.47 |
| *P* value | |
| 24-week-old AL Tg$^{Umod\text{wt}}$ vs 24-week-old AL Tg$^{Umod\text{C147W}}$ | *P*<0.001 |
| 24-week-old AL Tg$^{Umod\text{wt}}$ vs 48-week-old AL Tg$^{Umod\text{wt}}$ | ns |
| 24-week-old AL Tg$^{Umod\text{C147W}}$ vs 48-week-old AL Tg$^{Umod\text{C147W}}$ | *P*<0.0001 |
| 48-week-old AL Tg$^{Umod\text{wt}}$ vs 48-week-old AL Tg$^{Umod\text{C147W}}$ | *P*<0.0001 |
| 48-week-old AL Tg$^{Umod\text{wt}}$ vs 48-week-old CR Tg$^{Umod\text{C147W}}$ | *P*<0.01 |
| 48-week-old AL Tg$^{Umod\text{C147W}}$ vs 48-week-old CR Tg$^{Umod\text{C147W}}$ | *P*<0.0001 |
| 24-week-old AL Tg$^{Umod\text{C147W}}$ vs 48-week-old CR Tg$^{Umod\text{C147W}}$ | ns |

**Supplemental Table 7. Blood urea nitrogen and serum creatinine levels in 24-week-old AL Tg$^{Umod\text{wt}}$ and Tg$^{Umod\text{C147W}}$, 48-week-old AL Tg$^{Umod\text{wt}}$ and 48-week-old AL or CR Tg$^{Umod\text{C147W}}$ mice.** Data are mean ± s.d. n=8 mice/group. The *P* value for the specified comparisons (one-way ANOVA followed by Šídák’s correction) is reported. ns; not significant.

| **Diuresis (μl gBW$^{-1}$ h$^{-1}$)** | |
|---|---|
| Group | Mean ± s.d. |
| 24-week-old AL Tg$^{Umod\mathrm{wt}}$ | 1.80 ± 0.42 |
| 24-week-old AL Tg$^{Umod\mathrm{C147W}}$ | 3.90 ± 1.50 |
| 48-week-old AL Tg$^{Umod\mathrm{wt}}$ | 1.86 ± 0.08 |
| 48-week-old AL Tg$^{Umod\mathrm{C147W}}$ | 7.14 ± 1.98 |
| 48-week-old CR Tg$^{Umod\mathrm{C147W}}$ | 3.54 ± 1.50 |
| Normal range values[7] | 0.83 – 3.33 |
| *P* value | |
| 24-week-old AL Tg$^{Umod\mathrm{wt}}$ vs 24-week-old AL Tg$^{Umod\mathrm{C147W}}$ | $P$<0.05 |
| 24-week-old AL Tg$^{Umod\mathrm{wt}}$ vs 48-week-old AL Tg$^{Umod\mathrm{wt}}$ | ns |
| 24-week-old AL Tg$^{Umod\mathrm{C147W}}$ vs 48-week-old AL Tg$^{Umod\mathrm{C147W}}$ | $P$<0.001 |
| 48-week-old AL Tg$^{Umod\mathrm{wt}}$ vs 48-week-old AL Tg$^{Umod\mathrm{C147W}}$ | $P$<0.0001 |
| 48-week-old AL Tg$^{Umod\mathrm{wt}}$ vs 48-week-old CR Tg$^{Umod\mathrm{C147W}}$ | ns |
| 48-week-old AL Tg$^{Umod\mathrm{C147W}}$ vs 48-week-old CR Tg$^{Umod\mathrm{C147W}}$ | $P$<0.0001 |
| 24-week-old AL Tg$^{Umod\mathrm{C147W}}$ vs 48-week-old CR Tg$^{Umod\mathrm{C147W}}$ | ns |
| **Urinary creatinine (μmol gBW$^{-1}$ 24h$^{-1}$)** | |
| Group | Mean ± s.d. |
| 24-week-old AL Tg$^{Umod\mathrm{wt}}$ | 0.18 ± 0.04 |
| 24-week-old AL Tg$^{Umod\mathrm{C147W}}$ | 0.09 ± 0.01 |
| 48-week-old AL Tg$^{Umod\mathrm{wt}}$ | 0.14 ± 0.05 |
| 48-week-old AL Tg$^{Umod\mathrm{C147W}}$ | 0.05 ± 0.02 |
| 48-week-old CR Tg$^{Umod\mathrm{C147W}}$ | 0.14 ± 0.02 |
| Normal range values[5] | 0.11 – 0.25 |
| *P* value | |
| 24-week-old AL Tg$^{Umod\mathrm{wt}}$ vs 24-week-old AL Tg$^{Umod\mathrm{C147W}}$ | $P$<0.0001 |
| 24-week-old AL Tg$^{Umod\mathrm{wt}}$ vs 48-week-old AL Tg$^{Umod\mathrm{wt}}$ | $P$<0.05 |
| 24-week-old AL Tg$^{Umod\mathrm{C147W}}$ vs 48-week-old AL Tg$^{Umod\mathrm{C147W}}$ | $P$<0.05 |
| 48-week-old AL Tg$^{Umod\mathrm{wt}}$ vs 48-week-old AL Tg$^{Umod\mathrm{C147W}}$ | $P$<0.0001 |
| 48-week-old AL Tg$^{Umod\mathrm{wt}}$ vs 48-week-old CR Tg$^{Umod\mathrm{C147W}}$ | ns |
| 48-week-old AL Tg$^{Umod\mathrm{C147W}}$ vs 48-week-old CR Tg$^{Umod\mathrm{C147W}}$ | $P$<0.0001 |
| 24-week-old AL Tg$^{Umod\mathrm{C147W}}$ vs 48-week-old CR Tg$^{Umod\mathrm{C147W}}$ | ns |

**Supplemental Table 8. Diuresis and urinary creatinine levels in 24-week-old AL Tg$^{Umod\mathrm{wt}}$ and Tg$^{Umod\mathrm{C147W}}$, 48-week-old AL Tg$^{Umod\mathrm{wt}}$ and 48-week-old AL or CR Tg$^{Umod\mathrm{C147W}}$ mice.** Data are mean ± s.d. n=8 mice/group. The *P* value for the specified comparisons (one-way ANOVA followed Šídák's correction) is reported. ns; not significant.

| Gene | Experimental group | Fold change relative to 24-week-old AL Tg$^{Umodwt}$ | $Log_{10}$ (adj *P* value) relative to 24-week-old AL Tg$^{Umodwt}$ |
|---|---|---|---|
| *Ptprc* | 24-week-old AL Tg$^{UmodC147W}$ | 1.94 ± 0.22 | -1.95 |
| | 48-week-old AL Tg$^{Umodwt}$ | 1.27 ± 0.30 | -0.12 |
| | 48-week-old AL Tg$^{UmodC147W}$ | 4.05 ± 0.95 | -4.00 |
| | 48-week-old CR Tg$^{UmodC147W}$ | 1.77 ± 0.58 | -1.42 |
| *Cd68* | 24-week-old AL Tg$^{UmodC147W}$ | 8.69 ± 4.96 | -2.49 |
| | 48-week-old AL Tg$^{Umodwt}$ | 2.83 ± 1.27 | -0.11 |
| | 48-week-old AL Tg$^{UmodC147W}$ | 20.63 ± 4.18 | -4.00 |
| | 48-week-old CR Tg$^{UmodC147W}$ | 10.64 ± 5.75 | -3.70 |
| *Fut4* | 24-week-old AL Tg$^{UmodC147W}$ | 4.36 ± 2.06 | -2.17 |
| | 48-week-old AL Tg$^{Umodwt}$ | 2.53 ± 0.94 | -0.48 |
| | 48-week-old AL Tg$^{UmodC147W}$ | 9.46 ± 2.82 | -4.00 |
| | 48-week-old CR Tg$^{UmodC147W}$ | 3.25 ± 1.95 | -1.09 |
| *Cd19* | 24-week-old AL Tg$^{UmodC147W}$ | 4.13 ± 1.63 | -1.30 |
| | 48-week-old AL Tg$^{Umodwt}$ | 2.47 ± 1.25 | -0.10 |
| | 48-week-old AL Tg$^{UmodC147W}$ | 15.04 ± 6.52 | -4.00 |
| | 48-week-old CR Tg$^{UmodC147W}$ | 4.41 ± 2.26 | -1.40 |
| *Cd5* | 24-week-old AL Tg$^{UmodC147W}$ | 3.61 ± 1.41 | -1.35 |
| | 48-week-old AL Tg$^{Umodwt}$ | 1.90 ± 1.21 | -0.01 |
| | 48-week-old AL Tg$^{UmodC147W}$ | 11.99 ± 8.18 | -4.00 |
| | 48-week-old CR Tg$^{UmodC147W}$ | 3.57 ± 1.95 | -1.52 |
| *Vim1* | 24-week-old AL Tg$^{UmodC147W}$ | 4.47 ± 1.16 | -1.70 |
| | 48-week-old AL Tg$^{Umodwt}$ | 1.57 ± 1.24 | -0.01 |
| | 48-week-old AL Tg$^{UmodC147W}$ | 12.80 ± 4.88 | -4.00 |
| | 48-week-old CR Tg$^{UmodC147W}$ | 4.92 ± 1.66 | -1.82 |
| *Col1a1* | 24-week-old AL Tg$^{UmodC147W}$ | 3.84 ± 1.25 | -1.73 |
| | 48-week-old AL Tg$^{Umodwt}$ | 1.57 ± 1.01 | -0.03 |
| | 48-week-old AL Tg$^{UmodC147W}$ | 7.55 ± 3.18 | -4.00 |
| | 48-week-old CR Tg$^{UmodC147W}$ | 2.91 ± 1.55 | -0.85 |
| *Tgfb1* | 24-week-old AL Tg$^{UmodC147W}$ | 6.83 ± 3.32 | -2.15 |
| | 48-week-old AL Tg$^{Umodwt}$ | 1.29 ± 0.66 | 0.00 |
| | 48-week-old AL Tg$^{UmodC147W}$ | 22.22 ± 9.09 | -4.00 |
| | 48-week-old CR Tg$^{UmodC147W}$ | 4.86 ± 2.35 | -2.52 |
| *Acta2* | 24-week-old AL Tg$^{UmodC147W}$ | 3.56 ± 1.50 | -1.86 |
| | 48-week-old AL Tg$^{Umodwt}$ | 2.09 ± 0.69 | -0.12 |
| | 48-week-old AL Tg$^{UmodC147W}$ | 13.44 ± 4.88 | -4.00 |
| | 48-week-old CR Tg$^{UmodC147W}$ | 5.50 ± 1.16 | -2.55 |
| *Lcn2* | 24-week-old AL Tg$^{UmodC147W}$ | 14.77 ± 6.18 | -1.38 |
| | 48-week-old AL Tg$^{Umodwt}$ | 1.26 ± 0.85 | -0.0001 |
| | 48-week-old AL Tg$^{UmodC147W}$ | 40.04 ± 9.25 | -4.00 |
| | 48-week-old CR Tg$^{UmodC147W}$ | 16.53 ± 5.93 | -1.34 |
| *Havcr1* | 24-week-old AL Tg$^{UmodC147W}$ | 5.63 ± 1.30 | -3.05 |
| | 48-week-old AL Tg$^{Umodwt}$ | 1.83 ± 1.11 | -0.07 |
| | 48-week-old AL Tg$^{UmodC147W}$ | 11.51 ± 3.70 | -4.00 |
| | 48-week-old CR Tg$^{UmodC147W}$ | 5.68 ± 2.12 | -3.30 |

**Supplemental Table 9. Data used to generate the categorical bubble plot in Figure 5.** Fold change (mean ± s.d.) and $Log_{10}$ of the adjusted *P* value (one-way ANOVA followed by Dunnett's correction) relative to 24-week-old AL $Tg^{\mathit{Umod}\mathrm{wt}}$ of the kidney expression level of the specified genes in the indicated experimental groups. A value ≤-1.3 (corresponding to a *P* value ≤0.05) was considered statistically significant.

| Target gene | Primer forward (5' to 3') | Primer reverse (5' to 3') |
|---|---|---|
| *Acta2* | GCTACGAACTGCCTGACGG | GCTGTTATAGGTGGTTTCGTGGA |
| *Atl3* | GTATCAGTGGCTGGTGCTTTTCG | ATCAGAGCCTCCTCGCCATGAA |
| *Ccpg1* | CCACGAAGATGAGCTGGATGGT | CAGTAACGGTCCAACACCTCTC |
| *Ccl2* | TTCACCAGCAAGATGATCCCA | ACCTCTCTCTTGAGCTTGGTG |
| *Ccl5* | GTGCCCACGTCAAGGAGTATT | CCCACTTCTTCTCTGGGTTGG |
| *Ccl12* | CATCAGTCCTCAGGTATTGGCTGGA | CTTGGGGTCAGCACAGATCTCCTT |
| *Cd5* | CCCTTGCCAATTCGATGGGA | GGGCTGGAAATCAGAGCAGA |
| *Cd19* | ACCAGTTGGCAGGATGATGG | GCTGAGGAGCTGCATAGAGG |
| *Cd68* | CTGACAAGGGACACTTCGGG | AGGCCAATGATGAGAGGCAG |
| *Col1a1* | CTGACGCATGGCCAAGAAGA | ATACCTCGGGTTTCCACGTC |
| *Fam134b-1* | CTGCTCACCTTCCTGGGTG | TTCCCAGCTTTCAGTGAGGC |
| *Fam134b-2* | CATAATAGTCCACTCCTCGGCTTC | CTCAGTCTGGCTCTTTCATCTG |
| *Fut4* | GAGGTGGGTGTGGATGAACT | GTTGGATCGCTCCTGGAATA |
| *Hprt1* | CTGGTGAAAAGGACCTCTCGAAG | CCAGTTTCACTAATGACACAAACG |
| *Havcr1* | TTGGCATCTGCATCGCAGCCC | GGGAATGCACAACCGCTGCGT |
| *Lcn2* | TCCCCCTGCAGCCAGACTTCC | AGTAGCGACAGCCCTGGTCCTG |
| *Ptprc* | GTTCTGGGCTCCTTCCTCTT | GGAGACCAGGAAGTCTGTGC |
| *Retreg1* | AGACTGAGCCAGTGCATTGCAG | ACACTGCAGACCAGGAGGCAAA |
| *Retreg2* | CTCTGAGCTGTCAGATGAGGAG | TCTCGGTTCAGAGCCTCTTCTG |
| *Retreg3* | CTTCAGTGTCCGTGGCTACATG | CTGAGGACAGAAGGCAGCAAGT |
| *Rtn3* | CATAACCCTGACCCTACAACAC | GGAGCGTCTACTCTGTCCTTG |
| *Sec62* | TTCTACGTGCCAGAGGTGAAGG | GCGAACATCCAGGCTCACAGAA |
| *Sqstm1* | CGGCTTCCAGGCGCACTACC | CCACAGGCCCGTTGCAACCA |
| *Tex264* | GTGTGGATCTATGACCCAGTTCA | CACCTACTCTCATTTCTGCTGGC |
| *Tgfb1* | CCCGCGTGCTAATGGTGGACC | TGCACGGGACAGCAATGGGG |
| *Tg Umod* | ATGGACCAGTCCTGTCCTG | TCGTATGGGTATCCGGAGTT |
| *Total Umod* | ATGGACCAGTCCTGTCCTG | AGCAGCATCCAGGTCAAAGG |
| *Umod* R186S | TATGAGACCCTGACTGAGTACTGGC | TATGAGACCCTGACTGAGTACTGGA |
| *Vim1* | GGATCAGCTCACCAACGACA | GGTCAAGACGTGCCAGAGAA |

**Supplemental Table 10. List and sequences of primers used for gene expression analysis by SYBR Green real-time RT-qPCR**. *Tg Umod*, transgenic uromodulin.

| Target protein | Host | Dilution | Company | Code | Use |
|---|---|---|---|---|---|
| **Primary antibodies** | | | | | |
| Uromodulin | Sheep | 1:2000 | US Biological | T0850-02B | WB |
| | | 1:500 | | | IF |
| HA | Rat | 1:500 | Roche | #11867423001 | IF |
| HA | Rabbit | 1:1000 | Sigma | H6908 | WB |
| | | 1:500 | | | IF |
| Lipocalin-2 | Goat | 1:500 | R&D System | AF1857 | WB |
| | | 1:100 | | | IF |
| Vimentin | Rabbit | 1:1000 | Abcam | Ab92549 | IF |
| F4/80 | Rat | 1:100 | BioRad | MCA497GA | IF |
| p62 | Rabbit | 1:1000 | Sigma | P0067 | WB |
| | | 1:100 | | | IF |
| Lc3b | Rabbit | 1:100 | Cell signalling | #2775 | WB |
| β-actin | Mouse | 1:10000 | Sigma | A2228 | WB |
| Gapdh | Mouse | 1:1000 | Santa Cruz Biotechnology | sc-32233 | WB |
| Calreticulin | Rabbit | 1:1000 | Sigma | C4606 | IF |
| phospho-S6RP (Ser235/236) | Rabbit | 1:1000 | Cell Signalling | #2211 | WB |
| | | 1:1000 | | | IF |
| S6 Ribosomal Protein | Rabbit | 1:1000 | Cell Signalling | #2317 | WB |
| **HRP-conjugated secondary antibodies** | | | | | |
| Sheep IgG | Donkey | 1:7500 | Abcam | ab6900 | WB |
| Rabbit IgG | Donkey | 1:7500 | GE-Healthcare | NA934V | WB |
| Mouse IgG | Sheep | 1:7500 | GE-Healthcare | NA931V | WB |
| Goat IgG | Rabbit | 1:7500 | Agilent | P0449 | WB |
| **Fluorescent-conjugated secondary antibodies** | | | | | |
| Rabbit IgG 488 | Donkey | 1:500 | Thermofisher | A21206 | IF |
| Rabbit IgG 594 | Donkey | 1:500 | Thermofisher | A21207 | IF |
| Rat IgG 488 | Donkey | 1:500 | Thermofisher | A21208 | IF |
| Rat IgG 594 | Donkey | 1:500 | Thermofisher | A21209 | IF |
| Goat IgG 488 | Donkey | 1:500 | Thermofisher | A11055 | IF |

**Supplemental Table 11. List of antibodies used for Western blot and/or immunofluorescence analysis**. WB, western blot; IF, immunofluorescence.
